# ER Ca^2+^-levels control neuromodulator secretion by regulating L-type Ca^2+^-channel activity via its STIM1-interacting domain

**DOI:** 10.1101/2024.11.12.623153

**Authors:** Rein I. Hoogstraaten, Ruud F. Toonen, Matthijs Verhage

**Author notes:** To whom correspondence should be addressed: Matthijs Verhage, PhD or Ruud Toonen, PhD.

## Abstract

Regulated secretion is typically triggered by (local) increases in intracellular Ca^2+^, but the source of Ca^2+^, influx through voltage gated Ca^2+^ channels or release from the endoplasmic reticulum (ER), has distinct effects, particularly for neuropeptide secretion from dense-core vesicles (DCVs). Here, we show that in primary mouse neurons acute ER Ca^2+^ depletion by caffeine, cyclopiazonic acid or thapsigargin resulted in minute increases in bulk cytosolic free Ca^2+^ ([Ca^2+^]bulk) that did not trigger significant DCV exocytosis. Remarkably, following acute ER Ca^2+^ depletion, action potential (AP) trains triggered 50-90% less DCV exocytosis as compared to naïve neurons. In contrast, synaptic vesicle (SV) exocytosis was similar with/without acute ER Ca^2+^ depletion. Unexpectedly, acute ER Ca^2+^ depletion also reduced AP-induced [Ca^2+^]bulk-increases. L-type Ca^2+^-channel inhibitor nimodipine produced similar effects: reduced [Ca^2+^]bulk-increase, DCV-exocytosis but not SV exocytosis, i.e., for all three parameters a phenocopy of ER depletion. Finally, introducing L-type channels lacking STIM1 interaction sites restored DCV exocytosis following ER store depletion. We conclude that in mouse neurons, acute ER Ca^2+^ release is not effective in releasing neuromodulators. Instead, ER depletion activates a STIM1-dependent negative feedback loop that inhibits L-type Ca^2+^ channel activity, essential for DCV- but not SV-exocytosis.

## Introduction

Activity-dependent Ca^2+^ influx is essential for secretion in the two main secretory pathways in neurons: the release of classical neurotransmitters from synaptic vesicles (SVs), and the release of neuropeptides and neuromodulators (hereafter referred to as neuromodulators) from dense-core vesicles (DCVs). SV exocytosis depends on Ca^2+^ influx typically through P/Q-type and, to a lesser extent, N-type voltage gated Ca^2+^ channels (VGCC) in response single action potentials (Castillo et al., 1994; Luebke et al., 1993; Takahashi & Momiyama, 1993; Wheeler et al., 1994; Han et al., 2011; Kaeser et al., 2011; Südhof, 2013). In contrast, DCV exocytosis requires longer lasting depolarizations or high-frequency firing (Balkowiec & Katz, 2002; Lundberg et al., 1986; Persoon et al., 2018; Verhage et al., 1991; Xia et al., 2009) and it is unclear what sources of Ca^2+^ are relevant for neuronal DCV exocytosis, including which VGCCs.

Previous studies indicated the importance of L-type VGCCs for somatic and postsynaptic BDNF and NT3 release (Kolarow et al., 2007), while N-type channels were shown to be important for BDNF release from young hippocampal neurons (Balkowiec & Katz, 2002). In addition, several studies have implicated internal Ca^2+^ stores in neuromodulator secretion (Balkowiec & Katz, 2002; Blöchl & Thoenen, 1995; Canossa et al., 2001; Gärtner & Staiger, 2002; Griesbeck et al., 1999; Kolarow et al., 2007). At the Drosophila neuromuscular junction, ER Ca^2+^ release regulates DCV exocytosis in at least three ways. First, ER Ca^2+^ release is specifically required to mobilize DCVs (Shakiryanova et al., 2005). Second, ER Ca^2+^ release promotes capturing transiting DCVs after stimulation (Wong et al., 2009). Third, depleting the ER of Ca^2+^ prior to stimulation strongly reduced the release of neuromodulators from these boutons (Shakiryanova et al., 2011). In chromaffin cells the role of ER Ca^2+^ for secretory vesicle exocytosis is more modulatory: activity-induced Ca^2+^ influx is amplified by the ER via Ca^2+^ induced Ca^2+^ release (CICR), which contributes to DCV exocytosis (reviewed in García et al., 2006). Similarly, in hippocampal neurons, caffeine- or thapsigargin-induced Ca^2+^ release from the ER induces suboptimal (10-50%) neuromodulator release compared to high-frequency or high potassium stimulation (Balkowiec & Katz, 2002; Canossa et al., 2001; Griesbeck et al., 1999). However, ER Ca^2+^ depletion strongly reduced stimulation-induced neuromodulator secretion (Balkowiec & Katz, 2002; Canossa et al., 2001; Griesbeck et al., 1999; Kolarow et al., 2007). Together these studies suggest a complex role for ER Ca^2+^ release in DCV exocytosis in invertebrate motor neurons and a positive modulatory role in mammals with an unexplained, potentially indirect effect of ER Ca^2+^ depletion on the secretion capacity.

Mechanistic studies on how ER Ca^2+^ stores impact DCV exocytosis have produced contrasting results between species. For example, previous studies suggested that ER Ca^2+^ release triggers CaMKII activation as essential downstream effector for fusion (Kolarow et al., 2007; Shakiryanova et al., 2005; Wong et al., 2009), yet genetic deletion of CaMKII does not affect DCV exocytosis in mammalian neurons (Moro et al., 2020). In addition, the role of the ER during neuronal activity has also produced contrasting results, with some studies reporting ER Ca^2+^ release (Dittmer et al., 2017, 2019; Emptage et al., 2001; Llano et al., 2000; Unni et al., 2004) while others report ER Ca^2+^ buffering (de Juan-Sanz et al., 2017; Panzera et al., 2022).

To provide more mechanistic insight into how ER Ca^2+^ stores impact DCV exocytosis, we combined single vesicle resolution DCV fusion assays with direct ER and cytosolic Ca^2+^ imaging to investigate how the ER contributes to DCV exocytosis in adult mouse hippocampal neurons. In contrast to previous studies (Balkowiec & Katz, 2002; Canossa et al., 2001; Griesbeck et al., 1999), we find that acute ER Ca^2+^ release, in the absence of other stimuli, is insufficient to induce significant DCV exocytosis. We confirm previous findings that depleting ER Ca^2+^ strongly reduces activity dependent DCV fusion without affecting SV fusion. We discovered that this inhibition is not due to a direct role of ER Ca^2+^, but that ER Ca^2+^ depletion reduces DCV exocytosis by activating a STIM1 feedback loop onto L-type Ca^2+^ channels. Together, these findings suggest a new and important role for the ER in regulated secretion in which [Ca^2+^]ER status and ER Ca^2+^ depletion indirectly influence neuromodulator secretion by controlling L-type channel activity essential for DCV fusion.

## Results

### Depletion of ER Ca^2+^ diminishes DCV exocytosis

To address the role of ER Ca^2+^ dynamics in neuromodulator secretion, we quantified DCV fusion events before, during and after pharmacologically releasing Ca^2+^ from the ER. To release Ca^2+^ from the ER, we used caffeine which activates ryanodine receptors (RyR) which rapidly releases Ca^2+^ from the ER (Martín & Buño, 2003; Mcpherson et al., 1991), cyclopiazonic acid (CPA) and thapsigargin that block the sacro/endoplasmic reticulum Ca^2+^ ATPases (SERCAs), which in turn results in a passive leakage of ER Ca^2+^ (Camello et al., 2002)(See Figure S1A-C). Lentiviral expression of the DCV fusion reporter NPY-pHluorin (Figure 1A) was used to quantify fusion events in single, hippocampal neurons grown on glial micro islands, upon ER Ca^2+^ release and/or bursts of high frequency stimulation (8 x 50 action potentials (AP) at 50Hz interspaced by 0.5 seconds) (Farina et al., 2015a; Moro et al., 2020, 2021; Persoon et al., 2018, 2019; Puntman et al., 2021). Multiple bursts of high frequency stimulation (50Hz) are known to elicit the most efficient DCV fusion in single neurons (Persoon et al., 2018), resulting in 64.5 ± 1127 (median ± range) DCV fusion events during the first 8 bursts of 50 APs at 50Hz (Control = 125 ± 280; Caffeine = 97.5 ± 717; CPA = 48 ± 902; Thapsigargin = 36 ± 1126, ns *p* = 0.6297, See also Table S1 for detailed statistics). Interestingly, we did not observe robust DCV exocytosis during the 3-minute period of active (Caffeine) or passive (CPA, Thapsigargin) Ca^2+^ release from the ER, except in a few neurons (Figure 1B-D, Figure S1D). Finally, a second stimulation burst (8 x 50 APs at 50Hz) 3 minutes after ER depletion showed a 50 to 90% reduction in DCV fusion events in all neurons (Figure 1E, F). We conclude that Ca^2+^ release from the ER, in the absence of other stimuli, is not sufficient to induce DCV fusion. In addition, ER depletion strongly diminishes activity dependent DCV exocytosis in almost all neurons.

**Figure 1:**
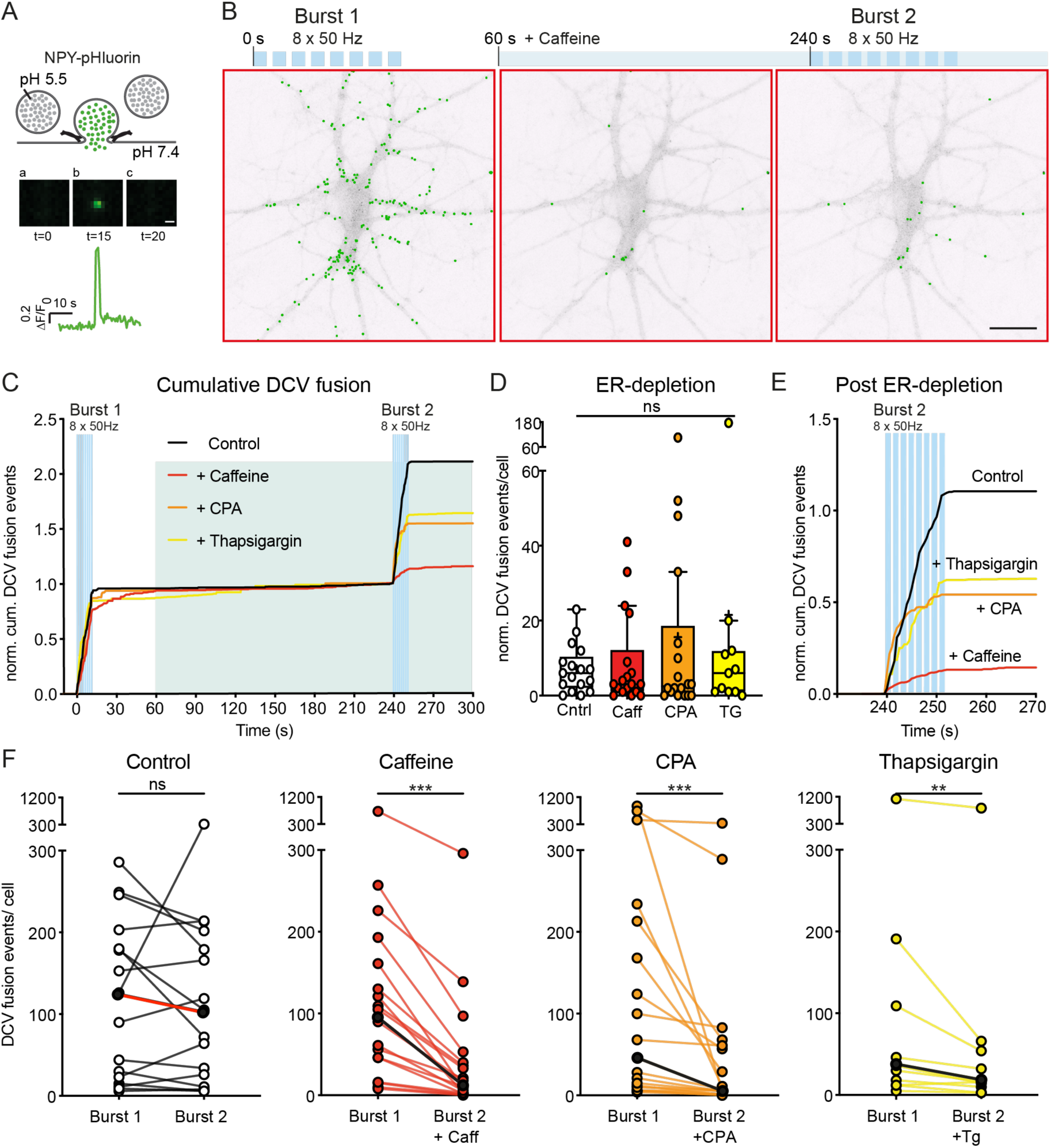
Depleting ER Ca^2+^ stores reduces neuronal DCV fusion. A) Schematic representation of DCV fusion assay using NPY-pHluorin. Before fusion, NPY- pHluorin is quenched in the acidic lumen of the DCV (a). During stimulation DCVs fuse with the plasma membrane visualized by a rapid increase in fluorescence (b) followed by a rapid decrease through cargo release or fusion pore closure and re-acidification (c). Scale bar 1 µm. Trace indicates F/F0 of a single DCV fusion event. B) Representative neuron with NPY-pHluorin labeled DCV fusion events superimposed in green showing fusion events during a burst of 8 x 50AP at 50Hz interspaced by 0.5 seconds (left) followed by superfusion of 20 mM caffeine (middle) and a subsequent burst 3 minutes after caffeine (right). upper bar indicates timing of stimulation (blue bars) and caffeine superfusion (light green). Scale bar, 25 µm. C) Cumulative median DCV fusion events over time normalized to the first burst. Blue bars depict stimulation, light green depicts superfusion with caffeine, CPA or thapsigargin. D) Boxplot of DCV fusion events in control or during the 3-minute treatment with caffeine, CPA or thapsigargin prior to burst 2. Boxplots represent median (line), mean (+) and Tukey range (whiskers). Each dot represents an individual neuron. Kruskal-Wallis with Dunn’s correction, ns = non-significant. E) Cumulative median DCV fusion events over time during burst 2. F) Number of DCV fusion events per neuron during the first and second stimulation (as in B) for control (N = 3, n = 17) or before and 3 minutes after treatment with 20 mM caffeine (N = 3, n= 18), 50 μM CPA (N = 3, n = 18) or 2 μM thapsigargin (N = 3, n = 11). Each individual neuron is connected with a line. The median during burst 1 and burst 2 are indicated with black dots connected with a red (control) or black line (rest). Wilcoxon matched-pairs signed rank test, ns = non-significant, **P < 0.01 ***P < 0.001.

### Depleting the ER Ca^2+^ affects cytosolic Ca^2+^ influx but not synaptic vesicle fusion

To explain the reduction in DCV exocytosis after depleting ER Ca^2+^, we tested how ER depletion affects cytosolic Ca^2+^ levels throughout a neuron. To this end, neurons were loaded with Fluo-5F to assess cytosolic Ca^2+^ in neurites (20 regions of interest, ROIs) during a train of action potentials, before and after depleting the ER using the same stimulation paradigm and pharmacological tools as in Figure 1. As shown before (Martín & Buño, 2003; Mcpherson et al., 1991), activation of the Ryanodine Receptor with caffeine acutely released Ca^2+^ from the ER into the cytosol, while blocking the SERCA pump with CPA resulted in a more gradual release of Ca^2+^ into the cytosol (Figure 2A, see also Figure S1).

**Figure 2:**
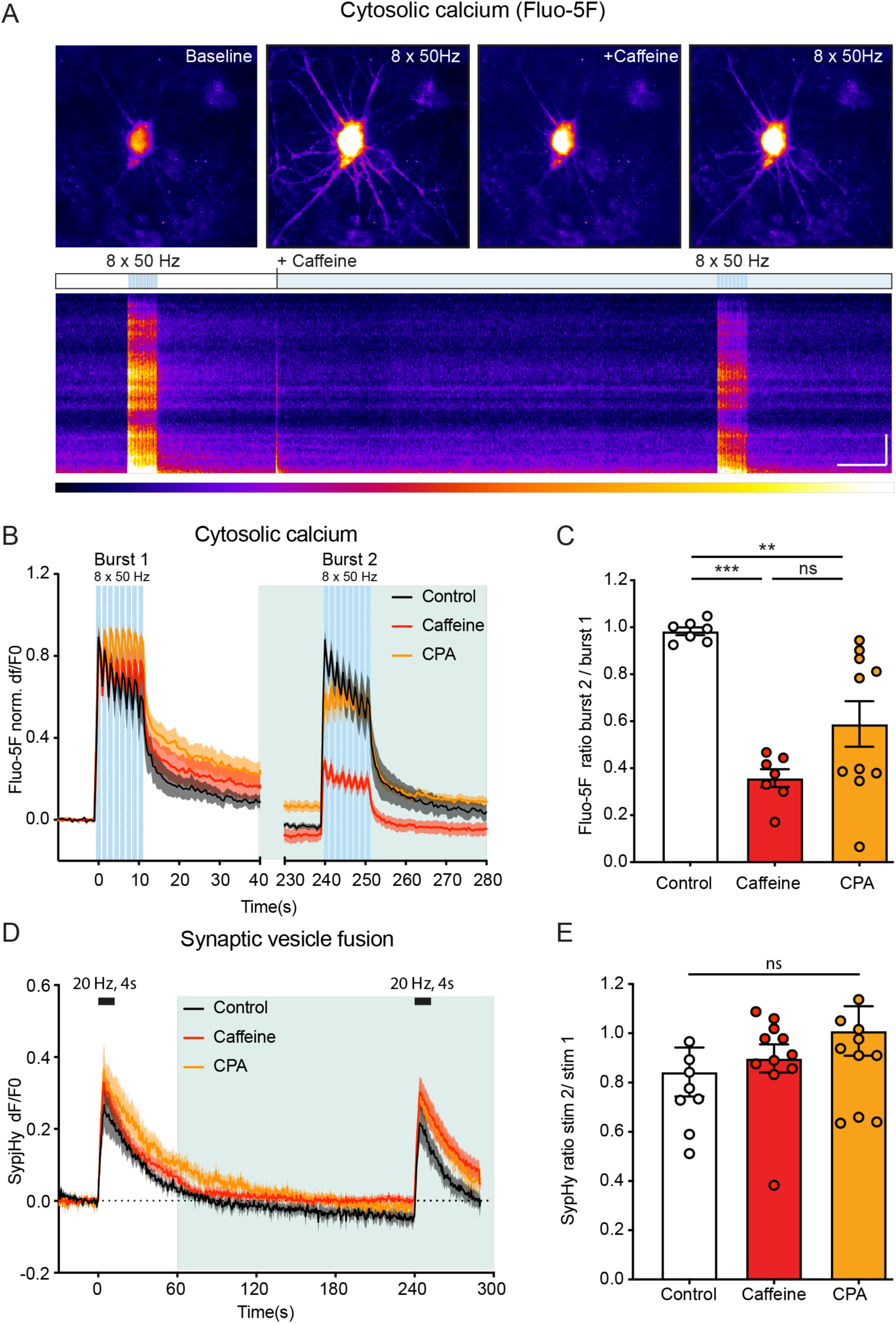
Depleting ER Ca^2+^ reduces activity induced cytosolic Ca^2+^ influx but not SV fusion. A) Typical example with kymographs of hippocampal neuron loaded with Fluo-5F stimulated with 8 x 50 AP at 50 Hz interspaced by 0.5 s (burst 1) followed by superfusion of 20 mM caffeine followed by another stimulation train (burst 2). Upper schematic indicates timing of stimulation (blue bars) and caffeine or CPA superfusion (light green). Pseudocolor scale shows low to high intensity from left to right. Scale bar, 10 µm, 20 seconds. B) Average dF/F0 traces over time of Fluo5-F responses during stimulation in control neurons (n = 7) or in neurons before and after caffeine (n = 7) or CPA (n = 11) treatment. C) ratio of Fluo-5F signal (area under the curve) between burst 2 and burst 1 for each condition. D) Average dF/F0 traces over time of SypHy responses during stimulation (20 Hz for 4s) in control neurons (n = 8) or in neurons before and after caffeine (n = 11) or CPA (n = 10) treatment. E) ratio of SypHy signal (max dF value) between stimulation 2 and stimulation 1 for each condition. Bars depict mean ± SEM. each dot represents an individual neuron. One-way ANOVA with Tukey’s (C) or Holm-Sidak (E) correction, ns = non-significant, ** p <0.01 *** p <0.001.

The total Ca^2+^ released from the ER was usually only observed after adding caffeine, while the more gradual release of Ca^2+^ after adding CPA could occasionally be observed (Figure 2A). However, the second train of action potentials, after ER Ca^2+^ depletion, resulted in a 40- 60% reduced Ca^2+^ influx compared to the first train (Figure 2A-C, Figure S1G). In conclusion, depleting ER Ca^2+^ strongly reduces subsequent action potential-induced Ca^2+^ influx.

To test whether ER depletion and the subsequent reduced Ca^2+^ influx affect synaptic vesicle exocytosis, we expressed synaptophysin-pHluorin as an SV fusion reporter and stimulated single hippocampal neurons with 80 AP at 20Hz. Unlike DCV exocytosis, ER depletion did not reduce stimulation-induced SV exocytosis compared to the same stimulation before ER depletion (Figure 2D-E). Together, these results show that ER Ca^2+^ depletion significantly reduces Ca^2+^ influx during stimulation without affecting SV exocytosis.

### The neuronal ER acts as Ca^2+^ source or sink during activity

Given the major impact of ER Ca^2+^ depletion on both Ca^2+^ influx and DCV exocytosis, together with conflicting findings regarding the role of the neuronal ER as a Ca^2+^ buffer (de Juan-Sanz et al., 2017; Panzera et al., 2022) or source (Dittmer et al., 2017, 2019) during neuronal activity, we next studied ER Ca^2+^ dynamics using ER-gCaMPMP6-150 (de Juan-Sanz et al., 2017). Administering a series of stimulations with a stepwise increase in frequency resulted in different responses between neurons, with some neurons showing ER Ca^2+^ uptake (buffering, Figure 3A-C, Neurons 1 and 2), while others showed ER Ca^2+^ release upon the same 20 Hz stimulation (Figure 3A-C, Neuron 3). ER Ca^2+^ dynamics varied over time, often changing from Ca^2+^ uptake to release after higher frequency stimulations (Figure 3A-C, neurons 1-3, from 50Hz onwards). Within different neuronal compartments (soma, synapses, axon, and dendrites), the ER responded uniformly at any point in time (Figure S2). These results indicate that the ER response is uniform at any point in time, as expected from a continuous organelle (Luarte et al., 2018; Terasaki, 2018; Wu et al., 2017; Yalçın et al., 2017), but functions as both a Ca^2+^ source or sink during activity and can change response direction over time.

**Figure 3:**
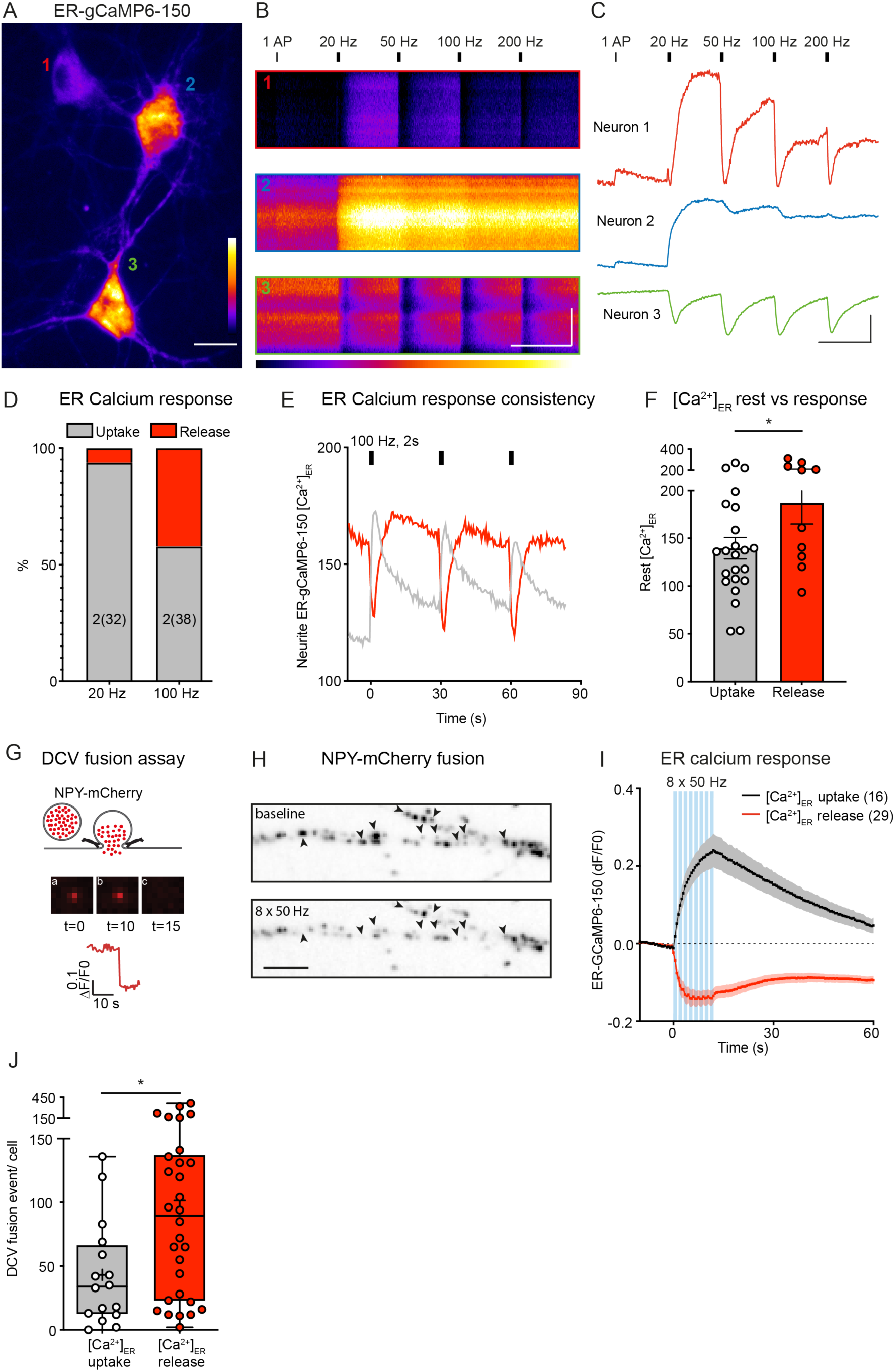
The ER acts as Ca^2+^ source or sink during neuronal activity. A) Hippocampal neurons infected with ER-GCaMP6-150 to visualize ER Ca^2+^. Image depicts signal of 3 neurons at baseline. The net ER-GCaMP6-150 signal in neuron 1 and 2 increases during the first AP and first 20 Hz stimulation indicating ER Ca^2+^ uptake, followed by ER Ca^2+^ release during the higher frequency stimulations whereas neuron 3 consistently releases Ca^2+^ already at 20 Hz stimulation. Neuron 1 even shows ER Ca^2+^ uptake, release and uptake within the 2 seconds of 20 Hz stimulation (Figure 3C). Pseudocolor scale shows low to high intensity from bottom to top. Scale bar, 20 µm. B) Kymograph of ER gCaMP6-150 response during stimulation of the 3 neurons in A. 1 action potential (AP) or 2 second stimulations with different frequencies are indicated on top. Pseudocolor scale below shows low to high intensity from left to right. Scale bar, 10 μm, 30 seconds. C) dF/F0 trace of the 3 neurons in A. Scale bar, 0.4 dF, 30 seconds. D) Percentage of neurons that take-up (gray bars) or release (red bars) ER Ca^2+^ (measured with ER-GCaMP6-150) during a 2 second 20 Hz or 100 Hz stimulation. Number of experiments and total number of neurons (within brackets) are indicated per condition. E) ER gCaMP6-150 response example of 2 neurons stimulated 3 X with 100 Hz (2 seconds). Y-axis indicates [Ca^2+^]ER concentration determined with ionomycin (see methods section). F) [Ca^2+^]ER concentration at baseline versus its response during stimulation. Bars depict mean ± SEM. Each dot represents an individual neuron. Unpaired t-test * p < 0.05. G) schematic representation of DCV fusion assay using NPY-mCherry. Before stimulation DCVs are visualized with NPY-mCherry (a). During stimulation DCVs fuse with the plasma membrane visualized by a rapid drop in fluorescence through cargo release (b and c and trace). Scale bar, 1 μm. H) Representative neurite stretch with NPY-mCh labeled DCVs before (upper) and during a burst of 8 x 50AP at 50Hz interspaced by 0.5 seconds (lower). Arrowheads indicate fusion events. Scale bar, 5 μm. I) Average trace of somatic ER-GCaMP6-150 response during bursts of 8 x 50AP at 50Hz interspaced by 0.5 seconds separated into neurons that take-up (black trace, N = 3, n = 16) or release (red trace, N = 3, n = 29) ER Ca^2+^. J) Boxplot of NPY-mCherry labeled DCV fusion events separated based on ER-GCaMP6-150 response. Boxplots represent median (line), mean (+) and Tukey range (whiskers). Each dot represents an individual neuron. Mann-Whitney U test, * p < 0.05.

Changes in ER Ca^2+^ concentration, due to previous activity, may influence the response at later time points. To determine whether stimulation frequency or [Ca^2+^]ER status prior to activity favors a certain ER response, we stimulated single neurons with either a low (20 Hz, 2 seconds) or high (100 Hz, 2 seconds) stimulation (Figure 3D-F). Furthermore, we determined the [Ca^2+^]ER at baseline by calibration with ionomycin to saturate the ER-GCaMP6-150 signal at the end of each experiment as previously described (de Juan-Sanz et al., 2017). At lower stimulation frequencies (20Hz), most neurons (94%) showed a net Ca^2+^ uptake into the ER (Figure 3D), in agreement with previous findings using the same stimulation frequency (de Juan-Sanz et al., 2017). On the contrary, at 100 Hz stimulation 42% of neurons showed acute ER Ca^2+^ release (Figure 3D). Neurons with a higher [Ca^2+^]ER at rest were significantly more likely to release ER Ca^2+^ during stimulation (Figure 3E-F). Furthermore, neurons that changed response over time most often did so when the ER concentration prior to the stimulation had changed compared to the initial baseline (Figure 3C and Figure S2F-H). Taken together, these results suggest that the ER’s dual role as Ca^2+^ source or sink during neuronal activity depends on its [Ca^2+^]ER status before stimulation and the strength of the stimulation, with higher stimulation frequencies and [Ca^2+^]ER favoring ER Ca^2+^ release.

### ER Ca^2+^ release correlates with increased DCV fusion

Given the strong dependence of DCV exocytosis on ER Ca^2+^ levels, but also the heterogenous response of the ER to neuronal activity, we next evaluated whether ER Ca^2+^ release correlates with increased DCV exocytosis. Therefore, we labeled neurons with ER-GCaMP6-150 and NPY-mCherry to simultaneously monitor ER Ca^2+^ dynamics and DCV exocytosis (Figure 3G- H). Neurons were stimulated with eight bursts of high frequency stimulation (8 x 50 action potentials at 50Hz interspaced by 0.5 seconds) and post hoc divided into 2 groups based on ER Ca^2+^ uptake or release (Figure 3I). On average, neurons that released ER Ca^2+^ during stimulation showed significantly more DCV exocytosis (Figure 3J). These results suggest that ER Ca^2+^ release contributes to DCV exocytosis and/or that high initial ER Ca^2+^ levels indirectly promote DCV exocytosis.

### Blocking L-type Ca^2+^ channels phenocopies ER Ca^2+^ depletion

Several studies have described a feedback loop that reduces Ca^2+^ influx upon ER depletion. This feedback loop is mediated by the Ca^2+^-dependent, ER-resident protein STIM1, which upon lowering of [Ca^2+^]ER, translocates to ER-PM contact sites and inhibits VGCCs, in particular, L-type Ca^2+^ channels (de Juan-Sanz et al., 2017; Dittmer et al., 2017; Park et al., 2010; Wang et al., 2010). Therefore, we hypothesized that blocking these channels phenocopies ER Ca^2+^ depletion.

Blocking L-type channels using nimodipine reduced volume average cytosolic Ca^2+^ influx in neurites and soma by 74% (Figure 4A-B). SV exocytosis remained unaffected (Figure 4C-D). On the contrary, blocking N- or P/Q- type channels did not affect cytosolic Ca^2+^ influx in neurites (Figure S3A-B), but blocking P/Q-type channels reduced SV exocytosis (Figure S3C- D), as expected. Since P/Q-type channels predominantly localize to the presynaptic active zone (Nakamura et al., 2015), we used syn-GCaMP6f to measure the effects of channel blockers more specifically on presynaptic Ca^2+^ influx. Blocking P/Q-type channels reduced Ca^2+^ influx at presynaptic release sites, but only upon low frequency (5 Hz) stimulation (Figure S3E-G). These results suggest that L-type channels mediate most of the neuronal Ca^2+^ influx in all neurites, and at higher frequency stimulation also in synapses.

**Figure 4:**
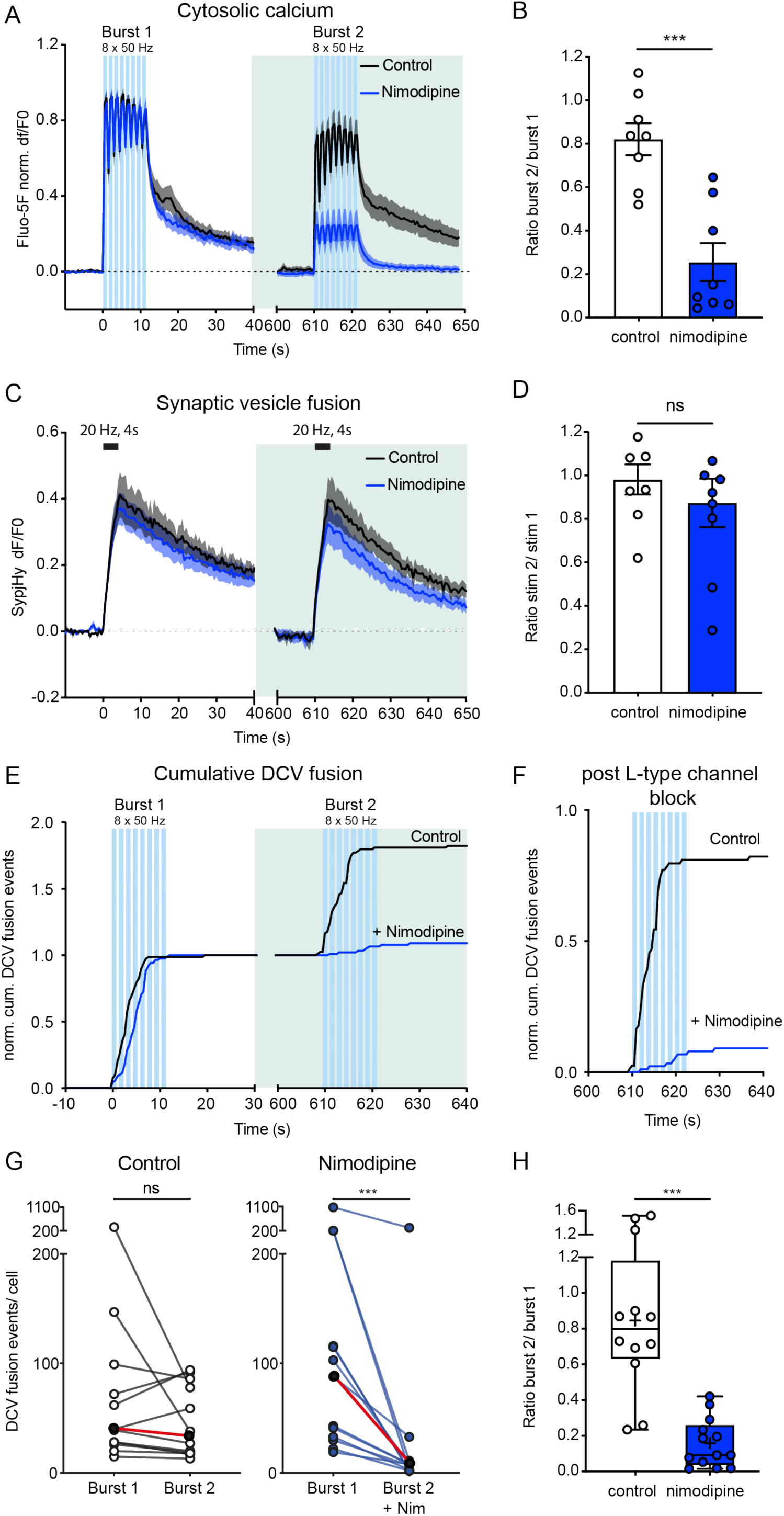
Blocking L-type voltage gated Ca^2+^ channels reduces DCV fusion. A) Average dF/F0 traces over time of Fluo5-F responses during stimulation in control neurons (N= 2 n = 8) or in neurons before and after nimodipine treatment (N = 2 n = 8). B) ratio of fluo-5F signal (area under the curve) between burst 2 and burst 1 for each condition. C) Average dF/F0 traces over time of SypHy responses during stimulation (20 Hz for 4s) in control neurons (n = 8) or in neurons before and after nimodipine treatment (n = 9). D) ratio of SypHy signal (max dF value) between stimulation 2 and stimulation 1 for each condition. Bars depict mean ± SEM. each dot represents an individual neuron. Unpaired t-test, ns = non-significant, *** p <0.001. E) Cumulative median DCV fusion events over time normalized to the first burst. Blue bars depict stimulation bursts, light green depicts superfusion with nimodipine. F) Cumulative median DCV fusion events over time during burst 2. G) Number of DCV fusion events per neuron stimulated twice with bursts of 8 x 50AP at 50Hz interspaced by 0.5 seconds, in control neurons (N = 3, n = 12) or before and 10 minutes after 30 μM nimodipine treatment (N = 3, n = 13) to block L-type channels. Each individual neuron is connected with a line. The median during burst 1 and burst 2 are indicated with black dots connected with a red line. Wilcoxon matched-pairs signed rank test, ns = non-significant, ***p < 0.001. H) Boxplot of the DCV fusion event ratio between burst 2 and burst 1 for each condition. Boxplots represent median (line), mean (+) and Tukey range (whiskers). Each dot represents an individual neuron. Unpaired t-test, *** p < 0.0001.

Blocking L-type Ca^2+^ channels strongly inhibited DCV exocytosis in all neurons (Figure 4E-G) with an average reduction of 85% (Figure 4H). Blocking P/Q- type channels reduced DCV fusion by 40% and blocking N-type channels had no significant effect (Figure S4A-E). Given the differential distribution of L-, P/Q- and N-type channels (Leitch et al., 2009; Nakamura et al., 2015; Tippens et al., 2008), we also determined the ratio of synaptic versus extrasynaptic fusion after blocking each VGCC (Figure S4F). On average 60% of all DCV fusion events occur at synapses (Figure S4G-H) as previously shown (Arora et al., 2017; Farina et al., 2015a; Persoon et al., 2019; van de Bospoort et al., 2012). Blocking P/Q or L-type, but not N-type channels reduced synaptic DCV fusion to a similar extent (Figure S4G-H). Taken together, these results suggest that L-type channels mediate most of the neuronal Ca^2+^ influx in neurites, and at higher frequency stimulation also in synapses, and are the most important VGCCs for neuronal DCV exocytosis. In addition, blocking L-type channels mimics depleting ER Ca^2+^ stores, suggesting that the ER Ca^2+^-depletion feeds back onto L-type channels.

### Removing the STIM1 binding domain of L-type channels disables the feedback loop activated by ER Ca^2+^ depletion

One way in which the ER and L-type Ca^2+^ channels may be functionally connected, is via the ER-resident protein STIM1. The STIM-ORAI activating region (SOAR, aa344-442) of STIM1 interacts with the C-terminal tail of the pore forming alpha1c subunit of Cav1.2 (aa1809-1908), blocking its activity (Park et al., 2010; Wang et al., 2010). Therefore, we designed constructs that expressed either the full subunit or a truncated alpha1c subunit (at aa 1809), lacking the STIM1 interaction site. Furthermore, we included an extracellular HA-tag (Altierab et al., 2002) and introduced a point mutation (T1037Y) that renders the channel resistant to nimodipine (Dolmetsch et al., 2001)(Figure 5A). Combining the expression of these constructs with nimodipine treatment creates a functional L-type channel knockin, removing all contribution of the endogenous L-type channels as described before (Dolmetsch et al., 2001). Given the size and the number of constructs required to constitute functional L-type channels, we performed these experiments in mass cultured neurons.

**Figure 5:**
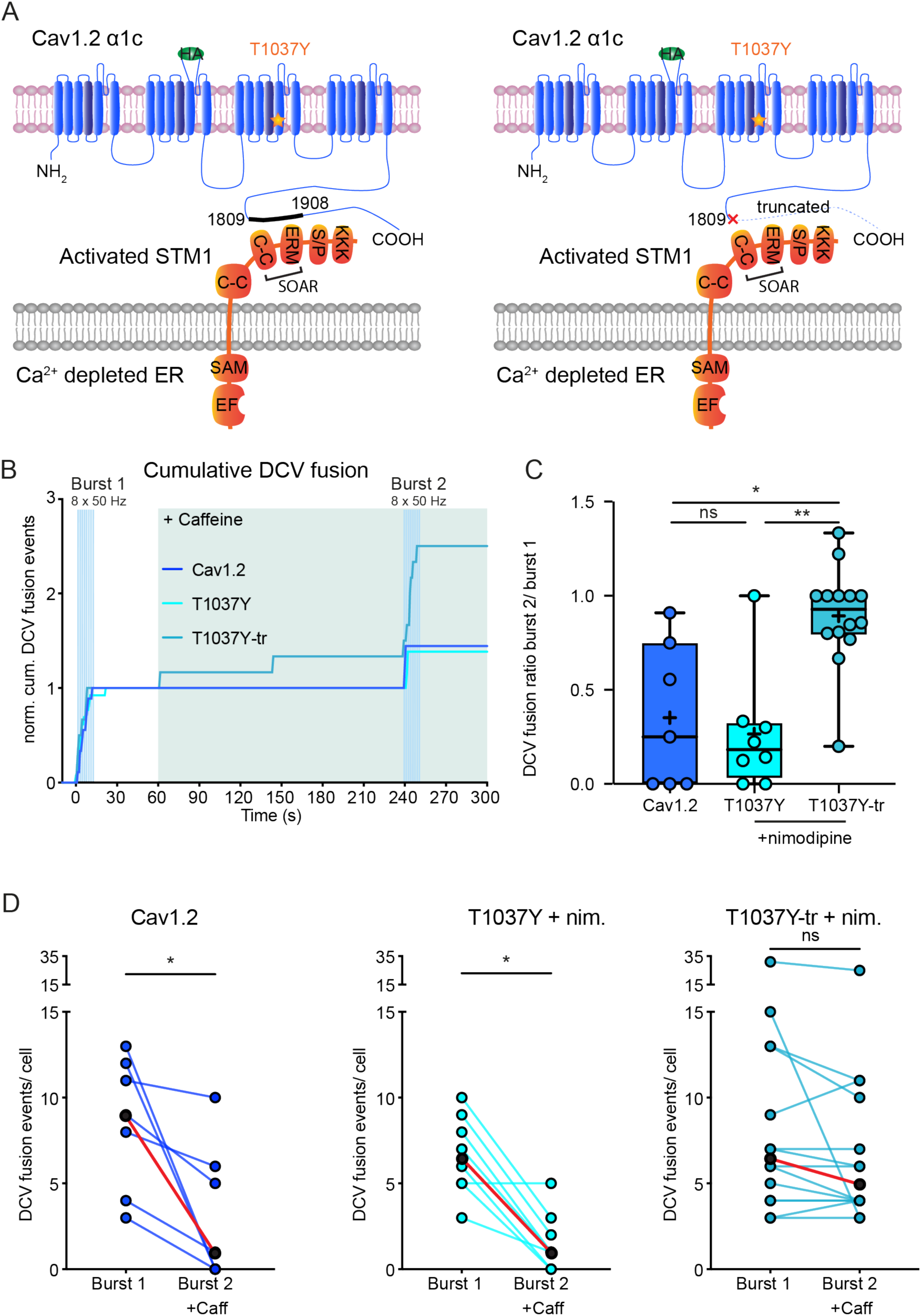
The C-terminal tail of L-type channels is essential for the ER Ca^2+^ depletion activated feedback loop that blocks DCV fusion. A) Schematic representation of L-type VGCC pore forming alpha-1c subunit including the location of the HA-tag, the T1037Y point mutation and the C-terminal tail of which aa1809- 1908 interact with the SOAR region of STIM1. Truncation of this C-terminal tail prevents STIM1 interaction and channel activation. B) Cumulative median DCV fusion events over time normalized to the first burst for the conditions in B. Blue bars depict stimulation bursts, light green depicts superfusion with caffeine (20 mM). C) Boxplot of the DCV fusion event ratio between burst 2 and burst 1 for each condition. Boxplots represent median (line), mean (+) and range (whiskers). Each dot represents an individual neuron. Kruskal-Wallis test with Dunn’s correction, ns = non-significant, * p<0.05, ** p < 0.01. D) Number of DCV fusion events per field of view stimulated twice with 8 x 50AP at 50Hz, once before (burst 1) and once 3 minutes after depleting ER Ca^2+^ with constant perfusion of 20mM caffeine (burst 2) in mass cultured hippocampal neurons transfected with Cav1.2 (N = 2, n = 7), T1037Y (N = 2, n = 8) or T1037Y truncated at aa 1809 (T1037Y-tr, N = 2 n = 14). All neurons were kept in 25 µM APV and DNQX containing Tyrodes. Neurons that expressed the nimodipine insensitive variant (TT1037Y and T1037Y-tr) were constantly kept in 5 µM nimodipine. Each individual neuron is connected with a line. The median during burst 1 and burst 2 are indicated with black dots connected with a red line. Wilcoxon matched-pairs signed rank test, ns = non-significant, ** p < 0.01, ***p < 0.001.

Neurons transfected with the alpha1c and the auxiliary b1b and a2d subunits showed co-immunostaining for Cav1.2 and the HA-tag (Figure S5A). In addition, HA-surface immunostaining labeled transfected neurons (Figure S5B), suggesting formation and membrane integration of mutant Cav1.2 channels.

The T1039Y point mutation (in rats, corresponding to T1037Y in mice) has been reported to reduce L-type channel sensitivity to nimodipine by 100-fold *in vitro* (He et al., 1997) and was used to create a functional knockin in several studies (Dolmetsch et al., 2001; Kamijo et al., 2018; Zhang et al., 2005, 2006). To test functional integration and nimodipine resistance of T1037Y containing constructs, we co-transfected all L-type channel subunits together with GCaMP6s to monitor Ca^2+^ influx, and stimulated neurons before and after incubation with 5 µM and subsequently 30 µM nimodipine (Figure S5C). 5 µM nimodipine reduced Ca^2+^ influx in all conditions (Figure S5C-D). However, this reduction was less severe in a substantial proportion of neurons expressing T1037Y and T1037Y-tr (Figure S5D). Treatment with 30 µM nimodipine abolished Ca^2+^ influx in all conditions (Figure S5E). These results confirm nimodipine resistance in T1037Y mutant expressing neurons at lower concentrations.

Next, we tested whether ER Ca^2+^ depletion still blocked DCV exocytosis in T1037Y and T1037Y-tr transfected neurons in 5 µM nimodipine. As an additional control we included Cav1.2 transfected neurons without nimodipine. The number of DCV fusion events in 1 field of view of hippocampal neurons grown in mass culture is generally lower compared to fusion in single neurons grown on glial micro islands (Figure 5B), as shown before (van de Bospoort et al., 2012). WT neurons that did not express heterologous L-type channels were excluded, as the number of fusing DCVs was too low to determine the relative effect of ER store depletion (Figure 5B).

Depleting ER Ca^2+^ with caffeine reduced DCV exocytosis in Cav1.2 expressing (without nimodipine) with 88% (Figure 5C-D). Likewise, DCV exocytosis in neurons expressing full length functional L-type mutant neurons (T1037Y) was reduced by 84% after ER Ca^2+^ depletion (Figure 5B-C). In contrast, in neurons expressing the truncated version without the STIM1 interaction site (T1037Y-tr), DCV exocytosis was not significantly reduced upon ER depletion (Figure 5B-C). These results suggest that the STIM1 interaction site is causally involved in the inhibition of DCV exocytosis by ER Ca^2+^ depletion.

## Discussion

In this study, we investigated different Ca^2+^ routes that regulate the release of neuromodulators from DCVs in hippocampal neurons at single vesicle resolution, especially the role of the neuronal ER as the primary intracellular Ca^2+^ store. Unexpectedly, prior depletion of ER Ca^2+^ substantially reduced DCV fusion events in response to action potential trains, whereas SV exocytosis was unaffected. In addition, blocking L-type channels phenocopied the effects of ER depletion. We identify the STIM1 binding domain in L-type channels as a causal link between ER Ca^2+^ levels and DCV exocytosis.

### Intact ER Ca^2+^ stores are essential for DCV fusion

Several studies have produced evidence to suggest that ER Ca^2+^ release can trigger neuromodulator release (Balkowiec & Katz, 2002; Blöchl & Thoenen, 1995; Canossa et al., 2001; Gärtner & Staiger, 2002; Griesbeck et al., 1999; Kolarow et al., 2007). Hence, the opposite conclusion as we reach here. These experiments measured bulk neuromodulator release from mass cultures after prolonged pharmacological ER depletion with low temporal resolution (Balkowiec & Katz, 2002; Blöchl & Thoenen, 1995; Canossa et al., 2001; Gärtner & Staiger, 2002; Griesbeck et al., 1999). Such extended ER calcium disruption likely elevates overall neuronal network activity in an indirect manner (Yoshimura, 2005). The increased network activity independently drives DCV secretion. Therefore, in the current study, we have prevented this non-specific effect by adding NMDA en AMPA blockers to the culture. Kolarow et al directly measured DCV exocytosis and, in agreement with our findings, observed reduced stimulation-induced DCV exocytosis after ER depletion. The authors argued that the simplest explanation for the reduced stimulation-induced exocytosis is that DCVs were already released during ER Ca^2+^ depletion (competition model, Kolarow et al., 2007). However, the current study shows that in hippocampal neurons the amount of calcium that the ER contributed to the overall increase of intracellular free Ca^2+^ in neurites was minute relative to Ca^2+^ influx through (mostly L-type) VGCCs and triggered no detectable DCV exocytosis (Figure 1); and that the reduced stimulation-induced exocytosis after ER Ca^2+^ depletion is due to an indirect effect depending on the STIM1 binding domain in L-type channels.

Instead of triggering neuromodulator release, we observed that ER Ca^2+^ store depletion leads to the opposite: Trains of action potentials that trigger efficient DCV exocytosis in naïve neurons, triggered substantially less after acute ER Ca^2+^ depletion in all neurons (Figure 1). The contribution of ER Ca^2+^ to DCV exocytosis is minute and cannot explain the strong effects after ER Ca^2+^ depletion based on a number of findings. First, ER Ca^2+^ release, in the absence of other stimuli, is insufficient to induce DCV fusion events (Figure 1). Second, the reduction in Ca^2+^ influx following ER depletion far exceeds the Ca^2+^ that the ER can provide as a source (Figure 2). Lastly, in a significant portion of neurons (approximately 35-60%), the ER operates primarily as a buffer for Ca^2+^ (Figure 3 and 4), and yet, depletion of the ER impacts fusion in all neurons. Therefore, we conclude that direct contributions of ER Ca^2+^ to the DCV fusion process are minimal and that ER Ca^2+^ depletion impacts DCV exocytosis in another, indirect manner.

### L-type channels conduct most neuronal Ca^2+^ influx, essential for DCV exocytosis

We find that L-type channel activity is essential the vast majority of neuronal DCV fusion events (Figure 4). The significance of L-type channels for BDNF release from neuronal somata and dendrites has been shown before (Kolarow et al., 2007; Xia et al., 2009). L-type channels are often only associated with somatic and postsynaptic calcium influx, even though ultrastructural evidence indicates axonal and occasionally presynaptic localization of these channels. In addition, a presynaptic form of LTP was found to depend on presynaptic BDNF release and L-type channel activity (Zakharenko et al., 2003). Our study extends beyond previous findings and suggest that L-type channels are essential for all DCV fusion events across the entire neuron, since the vast majority of DCVs (approximately 80%) fuse at axons (Persoon et al., 2018), and blocking L-type channels results in 85% reduction in fusion events (Figure 4).

Other studies found N-type channels to be more important for BDNF release, at least in young (DIV3) neurons (Balkowiec & Katz, 2002). We find no contribution to DCV exocytosis of N-type channels in more mature (DIV14-16) hippocampal neurons (Figure S3). SV exocytosis is known to transition from primarily using N-type channels to P/Q-type channels as they mature (Iwasaki et al., 2000; Scholz & Miller, 1995). A similar developmental shift from N-type to L-type channels may occur for DCV exocytosis.

Blocking P/Q type channels blocked 40% of DCV fusion events (Figure S3). Since blocking L-type channels blocked 85% (Figure 4), we conclude that substantial cooperativity exists between these channels. for DCV exocytosis. Given the enrichment of P/Q-type channels at the active zone (Nakamura t al., 2015), this cooperativity may mainly apply to DCV fusion events at presynaptic release sites. Indeed, blocking P/Q-type channels has a stronger impact on synaptic DCV fusion events and blocking L-type channels also inhibited synaptic DCV fusion events. This cooperativity between Ca^2+^ channels located at different locations in the neuron suggests that unlike SVs, DCVs are not coupled to specific channels at any subcellular location and their exocytosis, again unlike SVs, correlates well with volume average Ca^2+^ levels in neurites.

### STIM1 feedback onto L-type channels controls neuromodulator secretion

The current study demonstrates that the intracellular loop of the L-type Ca^2+^-channel, that binds STIM1, is causally involved in the way ER Ca^2+^ depletion impacts DCV exocytosis. Several previous studies have identified STIM1 as a link between ER Ca^2+^ depletion and L-type (Dittmer et al., 2017; Park et al., 2010; Wang et al., 2010) and potentially other VGCC (de Juan-Sanz et al., 2017). The current study shows that engineering L-type channels that lack the STIM1 interaction domain (Figure 5), mitigate the effects of ER Ca^2+^ depletion on DCV exocytosis. Therefore, we conclude that the ER controls neuromodulator secretion by regulating L-type channel activity via STIM1-based feedback. Our results establish a compelling new link between the ER Ca^2+^ levels and L-type channels that specifically controls neuromodulator exocytosis, without affecting SV exocytosis. However, STIM1 activation by ER store depletion also reduced single action potential-induced SV exocytosis (de Juan-Sanz et al., 2017), which suggests inhibition of other VGCCs, relevant for synaptic transmission, in addition to L-type channels. In the current study, using strong stimulation paradigms, we found no effect of ER Ca^2+^ depletion on SV exocytosis (Figure 2D-E).

### ER can act as Ca^2+^ source and sink

We find that the neuronal ER can act both as Ca^2+^ source and sink depending on the [Ca^2+^]ER status and stimulation frequency. Most studies have observed either one of these two (de Juan-Sanz et al., 2017; Dittmer et al., 2017, 2019; Emptage et al., 2001; Llano et al., 2000; Panzera et al., 2022; Unni et al., 2004). The fact that higher stimulation frequencies favor ER Ca^2+^ release may explain why some studies observed only Ca^2+^ buffering while using lower (20Hz -25Hz) stimulation (de Juan-Sanz et al., 2017; Panzera et al., 2022), while others found ER Ca^2+^ release only at high glutamate uncaging frequencies (Dittmer et al., 2017, 2019).

Studies on bullfrog sympathetic neurons and chromaffin cells also found higher [Ca^2+^]ER to favor release and vice versa (García et al., 2006; Verkhratsky, 2005), in line with our findings. These findings suggest that the neuronal ER Ca^2+^ buffering initially helps to secure low cytosolic Ca^2+^ levels, but higher influx rates saturate the ER Ca^2+^-ATPase, leading to RyR dependent Ca^2+^ release. Additionally, ER Ca^2+^ uptake rates slow down with increasing [Ca^2+^]ER (Verkhratsky, 2002), suggesting more saturated ER buffering capacities and more RyR dependent Ca^2+^ release when [Ca^2+^]ER at rest is higher.

We observed that ER-GCaMP6-150 fluorescence quickly returned to the same baseline value after each stimulation (Figure 3), with few exceptions (Figure S2), even though these baseline values often represented different [Ca^2+^]ER. This suggests homeostatic regulation of a certain resting [Ca^2+^]ER value that differs between neurons, regardless of prior neuronal activity.

## Limitations of the study

Monitoring DCV fusion after interference with the STIM1-L-type-channel interaction was technically challenging and required switching to transfection of multiple contrusts in mass cultured neurons. In addition, STIM1 has other important cellular functions than regulating Ca^2+^-channels, limiting the options for specific experimental manipulations to support its role in regulating DCV exocytosis. Despite these limitations, the relative rescue effects of truncated versus full length L-type channel clearly support our conclusion that ER Ca^2+^ depletion- suppresses activity-induced L-type channel Ca^2+^ influx, essential for DCV fusion, via its STIM1 interaction domain.

The implications of reduced neuromodulator secretion caused by this feedback loop are not yet clear. The notable magnitude of the observed inhibition implies its relevance, but the reduced neuromodulator secretion may still be a byproduct of a mechanism to reduce Ca^2+^ overload during ER stress, as suggested for STIM1 effects in dopamine neurons (Sun et al., 2017). Others suggested that the STIM1-dependent inhibition of synaptic transmission could be a way to reduce energy consumption during periods of ER stress (de Juan-Sanz et al., 2017). Such a mechanism may not be equally relevant for DCV fusion as these vesicles produce their own ATP via glycolysis (Hinckelmann et al., 2016). Yet reducing neuromodulator signaling, including all potential downstream pathways (GPCR/Trk signaling, receptor integration and transport, gene expression) could be an early coping strategy during ER stress. In contrast, reduced trophic factor signaling could actually aggravate persistent ER stress and even convey harmful effects to other cells. Finally, STIM1 feedback onto L-type channels also occurs at prolonged physiological stimulations that do not induce ER stress (Dittmer et al., 2017). This feedback loop could therefore directly limit DCV fusion under physiological conditions, in line with the observed DCV fusion rundown during intense stimulation (Persoon et al., 2018).

## Materials and Methods

### Animals

Animals were housed and bred in accordance with Dutch and institutional guidelines. All animal experiments were approved by the animal ethical committee of the VU University/VU University Medical Centre (license number: FGA 11-03 and AVD112002017824). Mice (C57Bl/6) mating was timed and E18 animals, acquired by caesarean section of pregnant mice, were used to dissect hippocampi or cortices.

### Neuron culture

Primary hippocampal neurons were cultured as described before (De Wit et al., 2009; Farina et al., 2015b). In short, dissected hippocampi were digested with 0.25% trypsin (Gibco) in Hanks’ balanced salt solution (Sigma) with 10 mM HEPES (Life Technologies) for 20 min. at 37 °C. Hippocampi were washed, triturated and counted prior to plating. For single hippocampal neurons, 1000-2000 neurons/well were plated on pre-grown micro-islands generated by plating 6000 rat glia on 18 mm glass coverslips coated with agarose and stamped with a solution of 0.1 mg/ml poly-D-lysine (Sigma) and 0.7 mg/ml rat tail collagen (BD Biosciences)(Mennerick et al., 1995; Wierda et al., 2007). For continental hippocampal cultures, 25.000 neurons/well were plated on pre-grown glial plates containing 18 mm glass coverslips. All neurons were kept in neurobasal supplemented with 2% B-27, 18 mM HEPES, 0.25% Glutamax and 0.1% Pen-Strep (all Gibco) at 37 °C and 5% CO_2_.

### Constructs

All constructs were cloned into lentiviral vectors containing a synapsin promoter to restrict expression to neurons. hSyn(pr)NPY-mCherry and hSyn(pr)NPY-pHluorin were used to quantify DCV fusion (Nagai et al., 2002; van de Bospoort et al., 2012). hSyn(pr)Synaptophysin-pHluorin was used to quantify SV fusion (Granseth et al., 2006). hSyn(pr)Synapsin-mCherry was used to label synapses (Persoon et al., 2019) and hSyn(pr)NavII-III to label axons (JoséGarrido et al., 2003). hSyn(pr)GCaMP6s was used for Ca^2+^ dynamics (T.-W. Chen et al., 2013) and hSyn(pr)ER-GCaMP6-150 was used for ER Ca^2+^ dynamics (de Juan-Sanz et al., 2017). pSyn(pr)cav1.2 (alpha1c subunit) was used together with constructs for the auxiliary subunits pSyn(pr)b1b (beta-1b subunit) and pSyn(pr)a2d (alpha-2-delta subunit) to express l-type channels. Nimodipine insensitive L-type channels were generated by substituting threonine 1037 for tyrosine (T1037Y) in the alpha1c subunit, similar to the rat T1039Y substitution (Dolmetsch et al., 2001). T1037Y-tr was truncated at aa 1809. All alpha1c subunits contained an HA tag between Q683 and T684, corresponding to the extracellular loop S5-H5 of domain II (Altierab et al., 2002).

### Lentiviral infections

For lentiviral work we followed all safety measures according to European legislation (ML-II, permit number: IG16-223-IIk). Lentiviral particles were produced as described before (Naldini et al., 1996). For DCV fusion experiments with NPY-pHluorin or NPY-mCherry and ER- GCaMP6-150 in island cultures, neurons were infected at DIV8-9 and imaged at DIV14-16. For ER Ca^2+^ dynamics with ER-GCaMP6-150 and Synapsin-mCherry or NAVII-III-mCherry island cultures were infected at DIV9 and imaged at DIV14-16.

### Calcium transfection

For calcium transfection of L-type channels we transfected the alpha1c (or T1037Y or T1037Y- tr), beta1b and alpha-2-delta at a 2:1:1 ratio. All cav1.2 subunits together were co-transfected with gCaMP6s or NPY-pHluorin at a 8:1 ratio. Neuronal cultures were pre-incubated with kynurenic acid (30 min - 1h at 37 °C) prior to transfection. For transfection we followed standard procedures to deliver plasmid containing calcium phosphate precipitates and incubated neurons for maximum 1h at 37 °C. Neurons were subsequently washed and kept at 37 °C until imaging. 25K continental cultures were transfected at DIV8 and imaged at DIV14- 15.

### Imaging

For confocal imaging of fixed samples imaging was performed on a A1R Nikon confocal microscope with LU4A laser unit (40x objective; NA 1.3) and NIS elements software (version 4.60, Nikon).

For life cell imaging experiments, coverslips were placed in an imaging chamber and imaged in Tyrode’s buffer (2 mM CaCl2, 2.5 mM KCl, 119 mM NaCl, 2 mM MgCl2, 20mM Glucose, 25mM HEPES, pH7.4) at RT. DCV or SV fusion experiments in single hippocampal neurons, were performed on a Zeiss AxioObserver.Z1 equipped with 561 nm and 488 nm lasers, a polychrome V, appropriate filter sets and AxioVision software (version 4.8, Zeiss) (Figures 1, 2, 4, 5, S1) or on a Olympus IX81 with an MT20 light source and appropriate filter sets (Semrock) and Xcellence RT imaging software (Olympus)(Figures S3, S4). Both setups contained a 40x oil objective (NA 1.3), an EMCCD camera (C9100-02; Hamamatsu, pixel size 200 nm). DCV fusion experiments in transfected continental cultures were performed on a Nikon Ti-E eclipse microscope with a LU4A laser system, appropriate filter sets, 40x oil objective (NA = 1.4) and EMCCD camera (Andor DU-897)(Figure 5). Continental cultures were imaged in Tyrode’s supplemented with 25 µM APV and 25 µM DNQX to prevent signal transmission after field stimulation. All DCV and SV pHluorin acquisitions were finalized with 5s of 25mM NH4^+^ perfusion dequenching pHluorin. For a detailed protocol of DCV fusion imaging and analyses see (Moro et al., 2021).

ER Ca^2+^ dynamics experiments were performed on a Nikon Ti-E eclipse microscope with a LU4A laser system, appropriate filter sets, 40x oil objective (NA = 1.4) and EMCCD camera (Andor DU-897) (Figures 3 and S2). Multiple positions were saved and revisited after incubation with 5 mM ionomycin (5 min at RT) to saturate the total ER-GCaMP5s signal used to calculate the [Ca^2+^]ER concentration.

All setups contained custom build perfusion barrels, and 2 parallel platinum electrodes that conducted 1 ms, 30 mA pulses delivered by a stimulus generator (A385RC) controlled by a Master 8 (AMPI). All images were acquired at 2Hz and all neurons were stimulated at indicated frequencies. For Ca^2+^ imaging with Fluo5-AM, neurons were incubated in 2 µM Fluo5-AM for 10 min at 37 °C. All other drugs were delivered by perfusion (Caffeine, CPA or thapsigargin) or by replacing tyrode’s and incubating cells (nimodipine, ω-agatoxin-IV, ω-conotoxin GVIIA, ionomycin). See table 2 for concentrations and incubation times.

| Name | final concentration | incubation time |
| --- | --- | --- |
| Caffeine | 20 mM | 3 minutes during acquisition |
| CPA | 50 $\mu\text{M}$ | 3 minutes during acquisition |
| Thapsigargin | 2 $\mu\text{M}$ | 3 minutes during acquisition |
| nimodipine | 30 $\mu\text{M}$ (Figures 4 and S5) 5 $\mu\text{M}$ (Figure 5 and S5) | 10 minutes at RT |
| ω-agatoxin-IV | 0.4 $\mu\text{M}$ | 10 minutes at RT |
| ω-conotoxin GVIIA | 3 $\mu\text{M}$ | 10 minutes at RT |
| ionomycin | 5 $\mu\text{M}$ | 5 minutes at RT |

### Immunocytochemistry

Neuronal cultures were fixed with 3.7% paraformaldehyde (PFA, Merck) in phosphate-buffered saline (PBS; 137 mM NaCl, 2.7 mM KCl, 10 mM Na_2_HPO_4_, 1.8 mM KH_2_PO_4_, pH 7.4) for 12 minutes at room temperature (RT). Cell were immediately immunostained or kept in PBS at 4 °C. Cells were permeabilized with PBS containing 0.5% Triton X-100 for 5 minutes and blocked with PBS containing 2% normal goat serum and 0.1% Triton X-100 for 1h at RT.

For surface staining of the cav1.2 HA-tag, cells were directly blocked with 2% normal goat serum in PBS. Cells were incubated with primary antibody overnight at 4 °C. Alexa fluor conjugated secondary antibodies (1:1000, Invitrogen) were incubated for 1 hour at RT. Coverslips were mounted in Mowiol (Sigma-Aldrich). Primary antibodies used were: polyclonal MAP2 (1:500, Abcam), polyclonal STIM1 (1:100, Cell Signaling), polyclonal STIM2 (1:100, Cell Signaling), monoclonal HA-tag (1:800, Covance), monoclonal Cav1.2 (1:100, Abcam) and monoclonal BDNF (1:10, DSHB).

### *de novo-*protein synthesis assay and Western blotting

For *de novo-*synthesized protein quantification, surface sensing of translation (SUnSET) was performed as described before (Schmidt et al., 2009). In brief, DIV 16-17 neurons were treated with 2 µM puromycin for 30 min at 37 before lysing. Lysates were run on a 10% SDS-PAGE gel containing 0.5% 2,2,2- trichloroethanol (TCE, Aldrich) and scanned with Gel Doc EZ imager (BIO-RAD) before transfer to measure total protein loading. Gels were subsequently transferred to a polyvinylidene difluoride membrane (Bio-Rad). Membranes were blocked with 5% milk (Merck) in PBS with 0.1% Tween-20 for 1 hour at RT and subsequently incubated with anti-puromycin (1:2500, Bio Connect) overnight at 4 °C. Allkaline phosphatase- conjugated secondary antibodies (1:10000, Jackson ImmunoResearch) were incubated for 50 minutes at RT. Puromycinylated proteins were subsequently visualized with AttoPhos (Promega) and scanned with an FLA-5000 fluorescent image analyzer (Fujifilm).

### Analysis

For DCV fusion experiments, 3x3 pixel ROIs were manually placed around each NPY-pHluorin fusion or NPY-mCherry event using ImageJ. A NPY-pHluorin event was considered a DCV fusion event if it suddenly appeared and if the maximal fluorescence was at least twice the SD above noise. A NPY-mCherry event was considered a fusion event if the signal suddenly dropped 2 SD below its initial intensity value. Custom written MATLAB (MathWorks, Inc.) scripts or ImageJ plugins (https://github.com/alemoro) were used to calculate the number and timing of fusion events. The synapsin-mCherry signal was used to determine synaptic versus extra-synaptic fusion. Custom written ImageJ plugins were used to segment synapsin-mCherry labeled synapses and to determine if the center of the DCV fusion ROI overlapped. For a detailed protocol of DCV fusion analyses see (Moro et al., 2021). For SV fusion experiments, 5x5 pixel ROIs were placed around 20 SypHy positive puncta per neuron to measure and calculate the average intensity over time per neuron using ImageJ and Excel. SV fusion was normalized to the NH4^+^ signal. For Ca^2+^ dynamics with Fluo5F-AM, gCaMP6s or ER-GCaMP6s-150, 5 pixel-wide lines were drawn on 5 neurites to calculate the average intensity over time per neuron using ImageJ and Excel. for ER Ca^2+^ dynamics in axons or dendrites, the axon was identified by the NavII-III signal. For (ER) Ca^2+^ dynamics in synapses, 5x5 pixel ROIs were placed around 20 synapsin-mCherry positive puncta per neuron to measure and calculate the average intensity over time per neuron. The ionomycin saturated intensity was used to calculate the [Ca^2+^]ER using the following formula from (de Juan-Sanz et al., 2017).

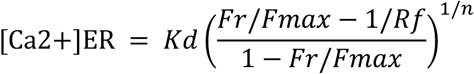

|  | Hill<br>coefficient<br>(n) | Rf<br>(Fsat/Fapo) | Kd (μM) | pKa (apo) | pKa (sat) |
| --- | --- | --- | --- | --- | --- |
| gCaMP6-150 | 1.6 | 45 | 150 | >12 | 7.09 |

### Statistics

Statistical tests were performed using R or GraphPad Prism. Normal distributions of all data were assessed with Shapiro-Wilk normality tests and Levene’s test of homogeneity of variances. To compare two groups, unpaired Student’s t-test in the case of normal distributed data or Mann-Whitney U tests for non-parametric data were used. For multiple comparisons one-way analysis of variance (ANOVA) followed by a post-hoc Tukey test to compare conditions was used when data was normally distributed or the Kruskal-Wallis test for non-parametric data followed by Dunn’s multiple comparisons test to compare conditions. Data is represented as boxplots (25-75% interquartile range) with the median (line), mean (+) and Tukey whiskers. See supplementary table 1 for detailed statistics.

## Acknowledgements

The authors would like to thank Joke Wortel and Desiree Schut for animal breeding and astrocyte culture and primary cell cultures assistance, Robbert Zalm for construct cloning and lentiviral production, Jian Dong for puromycin assay and Western blot, Alessandro Moro for analysis scripts and assistance and finally all members of the CNCR DCV team for fruitful discussions.

## Author contributions

R.I.H, R.T. and M.V. designed experiments. R.H. performed experiments and analyzed the data. R.I.H., R.T. and M.V. designed the figures and wrote the manuscript.

---

**Figure S1:**
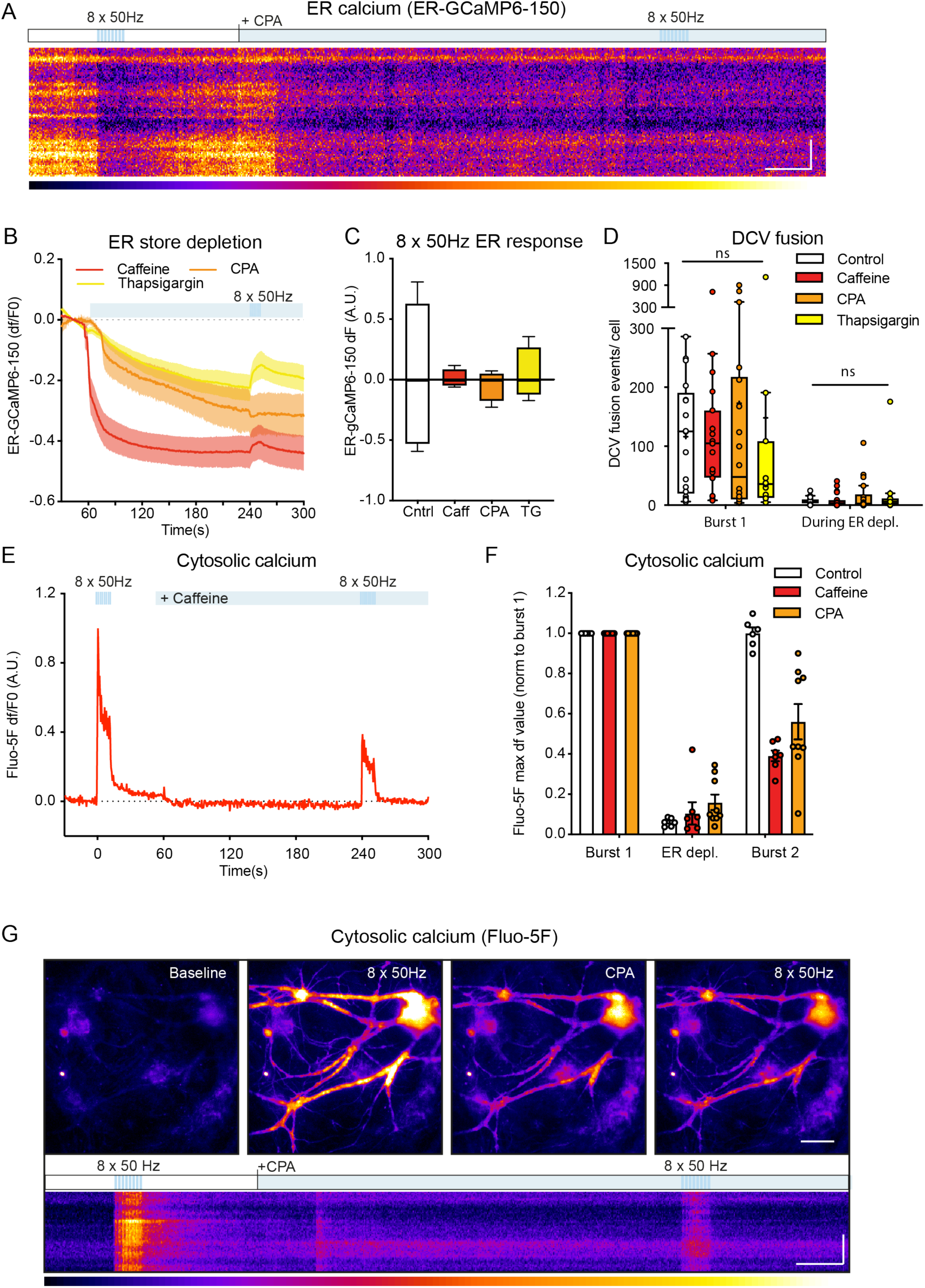
ER Ca^2+^ release does not induce DCV fusion. A) Kymograph of ER gCaMP6-150 response during ER store depletion with 50 μM CPA. Pseudocolor scale below shows low to high intensity from left to right. Scale 10 μm, 20 seconds. B) Average df/f0 trace of ER-GCaMP6-150 during treatment with caffeine (n = 24), CPA (n = 15) or thapsigargin (n = 7). C) Maximum or minimum ER-GCaMP6-150 response during the second 8 x 50 Hz stimulation for control neurons (n = 19) or neurons treated with caffeine (n = 24), CPA (n = 15) or thapsigargin (n = 7). D) Boxplot of DCV fusion events during a stimulation burst of 8 x 50 AP at 50 Hz interspaced by 0.5 s (burst 1) and during 3 minutes of perfusion with tyrodes (Control N = 3, n = 17) or 20 mM caffeine (N = 3, n = 18), 50 μM CPA (N = 3, n = 18) or 2 μM thapsigargin (N = 3, n = 11). Boxplots represent median (line), mean (+) and Tukey range (whisk- ers). Each dot represents an individual neuron. Kruskal-Wallis with Dunn’s correction, ns = non-significant. E) dF/F0 trace over time of Fluo-5F loaded neurite stretch corresponding to the kymograph in Figure 2A. F) max Fluo-5F response normalized to burst 1, during ER depletion and during burst 2 for each condition. G) Typical example with kymographs of a hippocampal neurite loaded with Fluo-5F stimulated with 8 x 50 AP at 50 Hz interspaced by 0.5 s (burst 1) followed by superfusion with 50 μM CPA followed by another stimulation train (burst 2). Pseudocolor scale shows low to high intensity from left to right. Scale bar, 10 μm, 20 seconds.

**Figure S2:**
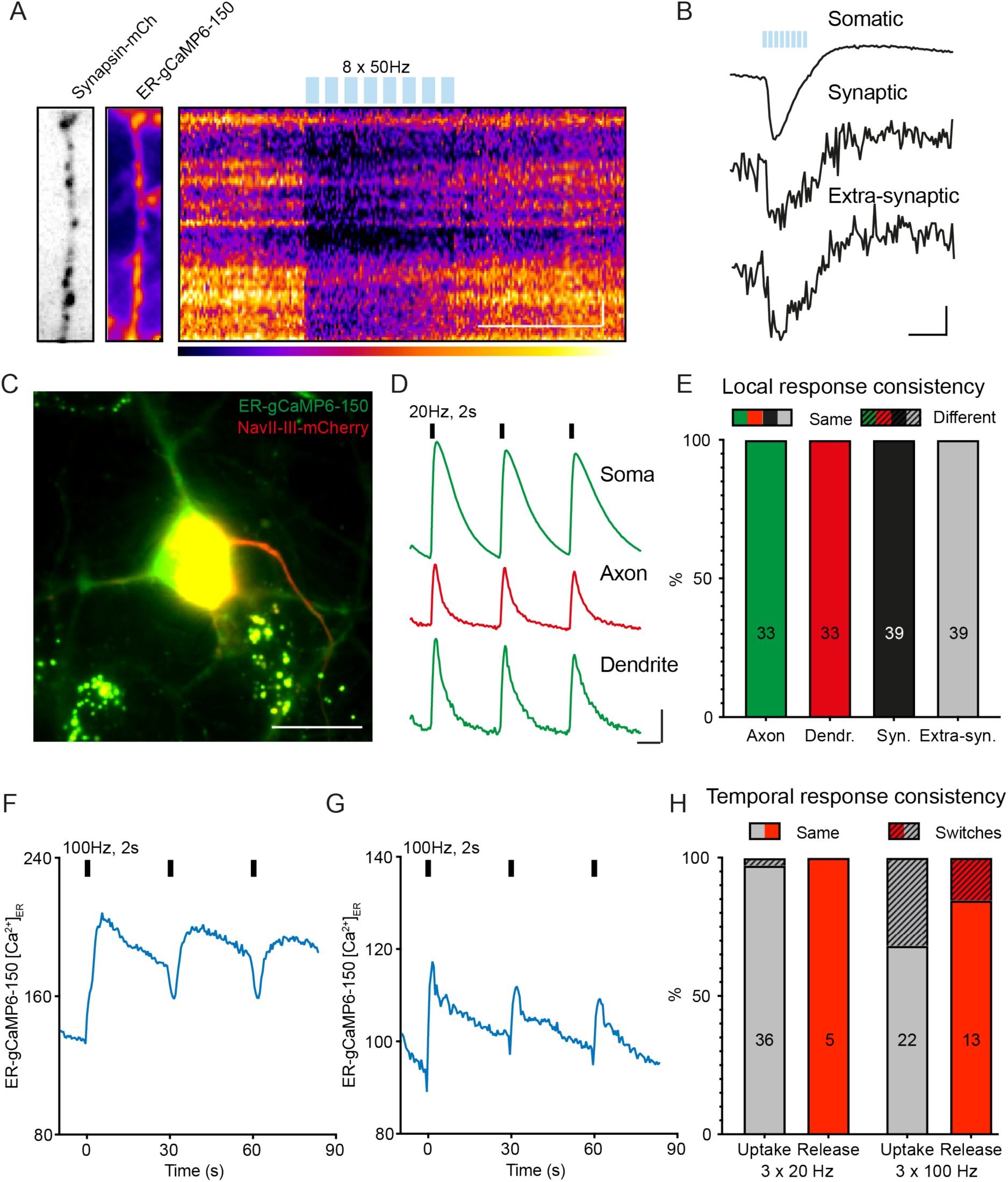
ER response to stimulation is consistent throughout each neuron. A) Hippocampal neurite expressing synapsin-mCherry and ER-GCaMP6-150. Kymograph shows ER-GCaMP response to a burst of 8 X 50 AP at 50 Hz interspaced by 0.5 seconds (blue bars). Pseudocolor scale shows low to high intensity from left to right. Scale bar, 5 μm, 10 seconds. B) ER-GCaMP6-150 dF/F0 traces of neuron in A in the soma, within or outside synapses. Blue bars indicate stimulation. Scale bar, 0.1 dF/F0, 10 seconds. C) Hippocampal neuron expressing ER-GCaMP6-150 (green) and NavII-III-mCherry to label the axon initial segment. Scale bar, 20 μm. D) ER-GCaMP6-150 dF/F0 traces of the neuron in C stimulated 3 times with a 20Hz stimulation (2 seconds). Response is shown for the soma, the NAVII-III labelled axon and the dendrites. Scale bar, 0.2 dF/F0, 10 seconds. E) Percentage of neurons that show the same or a different ER-GCaMP6-150 response to stimulation in different neuronal structures relative to somatic response of that same neuron. Total number of neurons are indicated per neuronal structure. F-G) ER gCaMP6-150 response examples of 2 neurons stimulated 3 X with 100 Hz (2 seconds) that switch between ER Ca^2+^ uptake and release over time. Y-axis indicates [Ca^2+^]ER concentration determined with ionomycin (see methods section). H) Percentage of neurons that switch ER-GCaMP6-150 response between ER Ca^2+^ uptake or release over time following 3 x 20 Hz stimulation (as in D) or 3 x 100 Hz stimulation (as in F and G). Total number of neurons are indicated per type of response.

**Figure S3:**
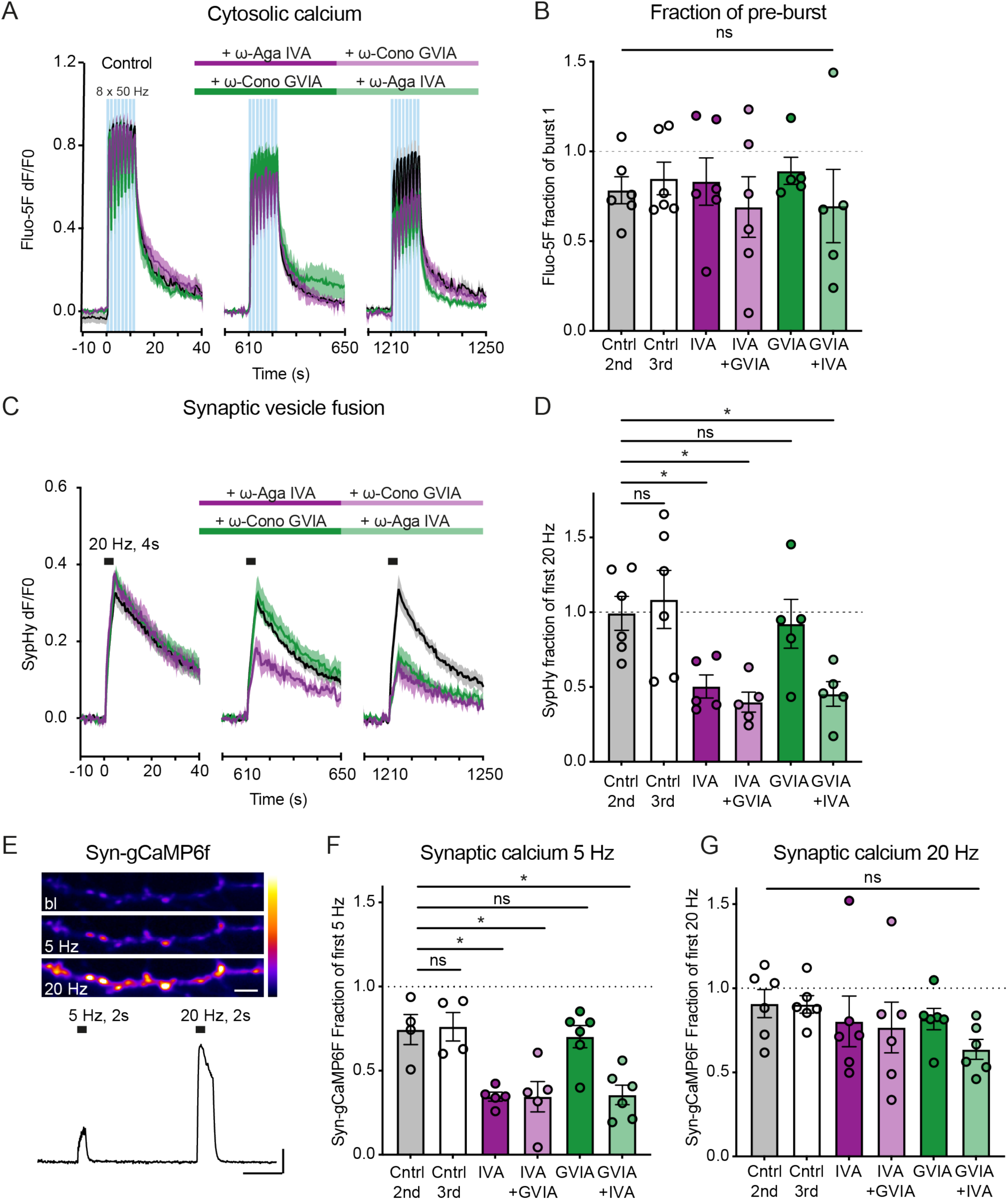
Blocking P/Q-type channels affect SV fusion but not bulk Ca^2+^ influx. A) Average dF/F0 trace over time of Fluo-5F responses during 3 bursts of 8 x 50AP at 50Hz interspaced by 0.5 seconds (10 minutes between bursts) in control neurons (N = 3, n = 6), neurons treated with ω -agatoxin IVA before burst 2 and ω -conotoxin GVIA before burst 3 (N = 3, n = 6), and neurons treated with ω -conotoxin before burst 2 and GVIA ω -agatoxin IVA before burst 3 (N = 3, n = 5). Blue bars depict stimulation bursts, purple and light purple indicate consecutive superfusion with ω -agatoxin IVA and ω -conotoxin GVIA respectively, green and light green indicate superfusion with ω -conotoxin GVIA and ω -agatoxin IVA respectively. B) Fluo-5F fraction of burst 1 (area under the curve) for burst 2 and burst 3 for each condition. C) Average dF/F0 traces over time of SypHy responses during stimulation (20 Hz for 4s) in control neurons (N = 3, n = 6), neurons treated with ω -agatoxin IVA before the second 20Hz and ω -conotoxin GVIA before the third 20Hz (N = 3, n = 5), or vice versa (N = 3, n = 5). D) Syphy fraction of the first 20Hz stimulation (max dF value) for the second and third 20Hz stimulation for each condition. E) Hippocampal neurite stretch expressing syn-gCaMP6f at baseline (bl), during 5Hz or 20Hz stimulation (2 seconds). Pseudocolor scale shows low to high intensity from bottom to top. Scale bar, 5 μm. trace depicts dF/F0 during both stimula- tions. Scale bar, 0.1dF/F0, 10 seconds. F) Syn-gCaMP6f fraction of the first 5Hz stimulation for the second and third 5Hz for each condition (control, n = 4, IVA-GVIA, n = 5, GVIA-IVA, n = 6). G) as in F but for 20Hz stimulation (control, n = 6, IVA-GVIA, n = 6, GVIA-IVA, n = 6). Bars depict mean ± SEM. each dot represents an individual neuron. One-way ANOVA with Tukey’s correction, ns = non-significant, * p <0.05.

**Figure S4:**
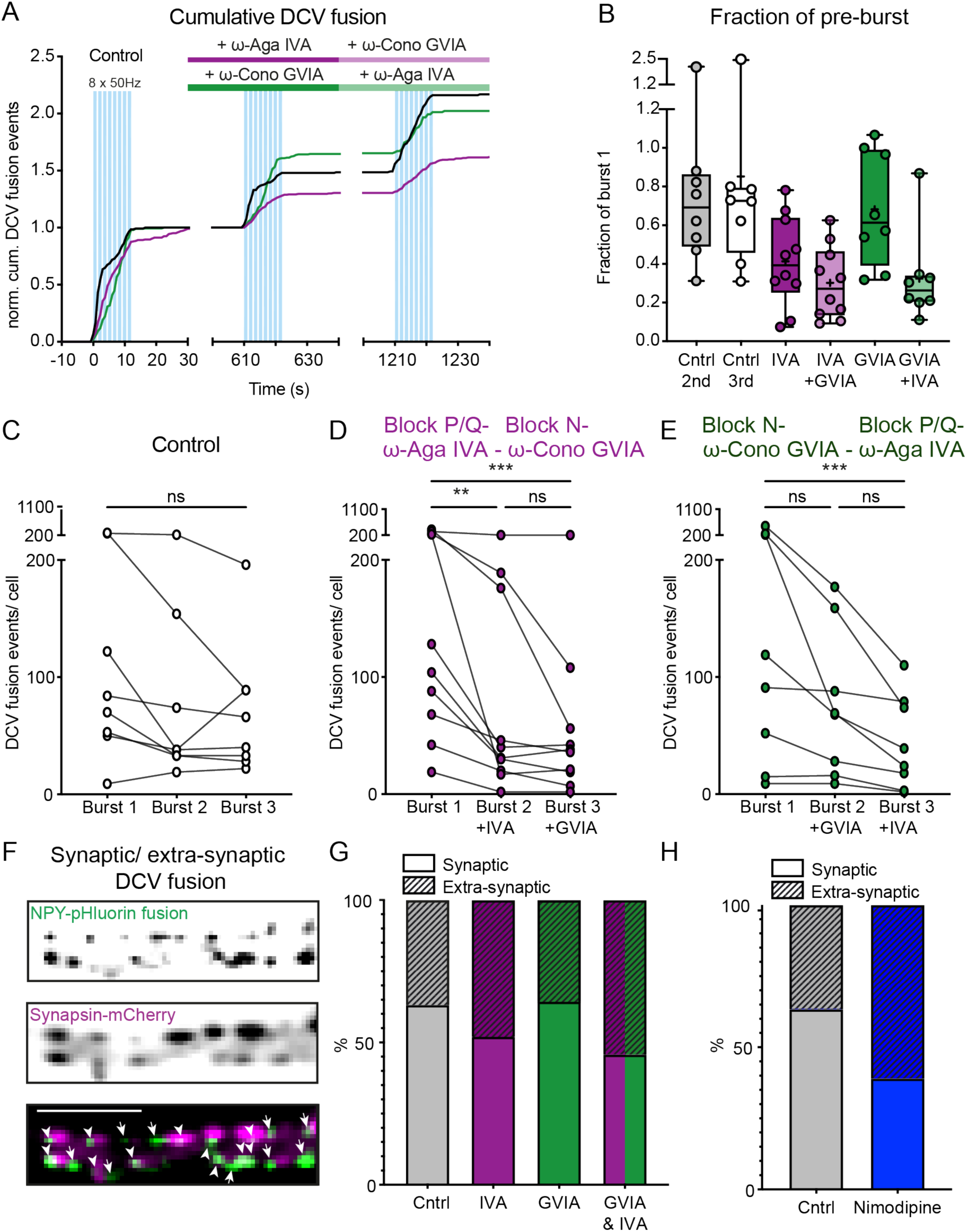
Blocking P/Q- type but not N-type voltage gated Ca^2+^ channels reduces DCV fusion. A) Cumulative median DCV fusion events over time normalized to the first burst. Blue bars depict stimulation bursts, purple and light purple indicate consecutive superfusion with ω - agatoxin IVA and ω -conotoxin GVIA respectively, green and light green indicate superfusion with ω -conotoxin GVIA and ω -agatoxin IVA respectively. B) Boxplot of the fraction of DCV fusion events during the second or third burst relative to the first burst per condition. Boxplots represent median (line), mean (+) and Tukey range (whiskers). Each dot represents an individual neuron. C) Number of DCV fusion events per neuron stimulated 3x with bursts of 8 x 50AP at 50Hz interspaced by 0.5 seconds (10 minutes between bursts) in control neurons (N = 3, n = 8). D) As in C but treated with ω -agatoxin IVA (10 minutes before burst 2) to block P/Q type channels and subsequently with ω-conotoxin GVIA (10 minutes before burst 3) to block N- type channels (N = 3, n = 10). E) Same as in C but treated with ω -conotoxin GVIA and subsequently with ω -agatoxin IVA (N = 3, n = 8). Each individual neuron is connected with a line. Friedman’s test with Dunn’s correction, ns = non-significant, **P < 0.01 ***P < 0.001. F) Representative neurite showing NPY-pHluorin labelled DCV fusion and synapsin-mCherry labeled synapses. Arrowheads indicate synaptic DCV fusion and arrows extra-synaptic DCV fusion. Scale bar, 5 μm. G) Percentage of synaptic and extra-synaptic DCV fusion per condition. The condition where both P/Q-type and N-type channels were blocked (burst 3 in B and C) are grouped together. H) Percentage of synaptic and extra-synaptic DCV fusion in control or after blocking L-type channels with nimodipine (related to Figure 4).

**Figure S5:**
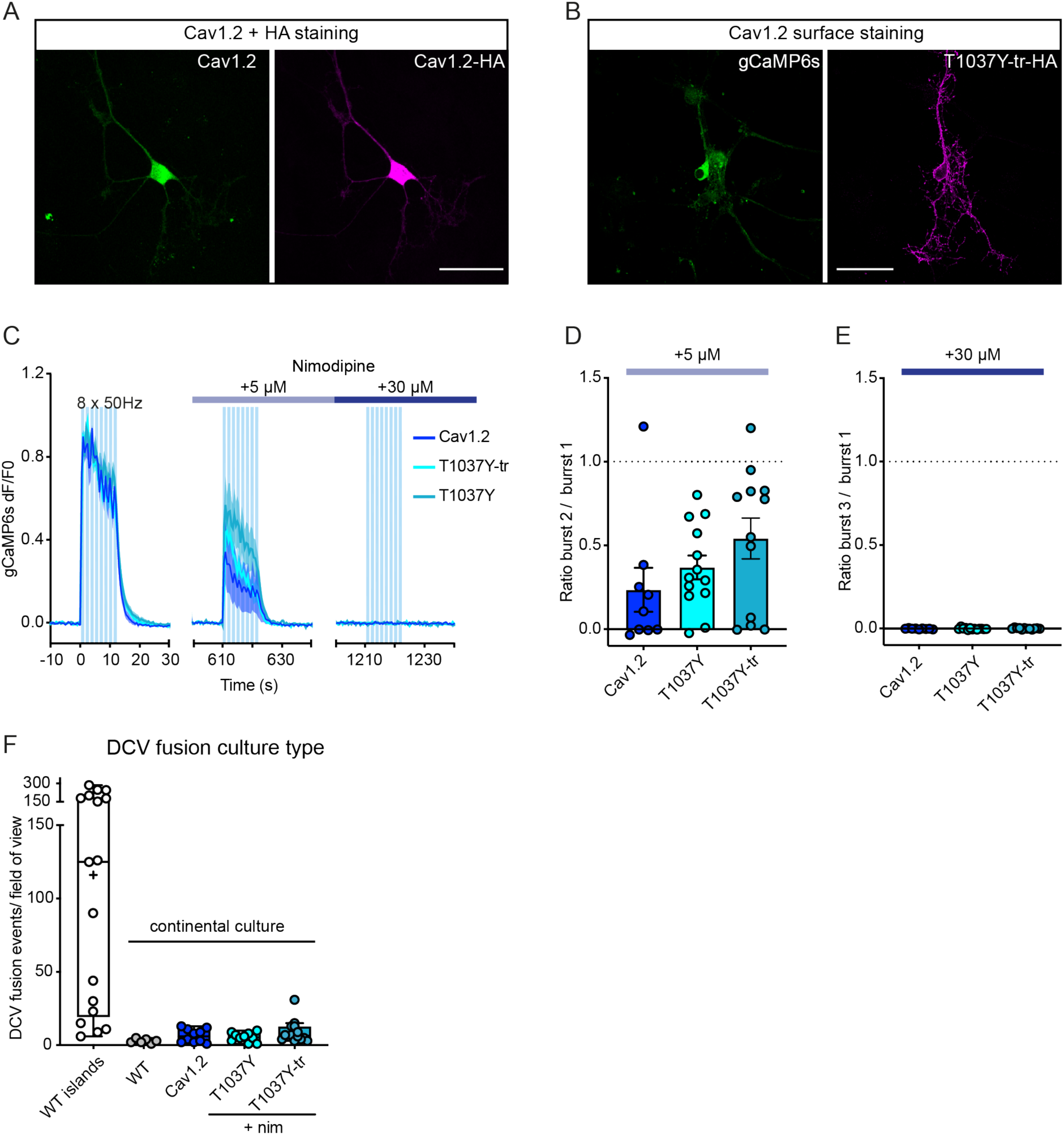
Functional L-type knockin verification. A) Neuron transfected with Cav1.2 immunostained for Cav1.2 (green) and the HA-tag (magenta). B) Neuron transfected with GCaMP6s (green) and T1037Y-tr surface immunostained (without permeabilization) for the HA-tag (magenta). Scale bars in A and B = 50 µm. C) Average dF/F0 trace over time of GCaMP6s responses during 3 bursts of 8 x 50AP at 50Hz interspaced by 0.5 seconds (10 minutes between bursts) of neurons transfected with full length Cav1.2 (n = 9), T1037Y (n = 13) or T1037Y-tr (n = 12). Blue bars depict bursts of 8 x 50AP at 50Hz interspaced by 0.5 seconds. All conditions were treated with 5 µM nimodipine 10 minutes before burst 2 (transparent dark blue bar) and 30 µM nimodipine 10 minutes before burst 3 (dark blue bar). All neurons were imaged in Tyrode’s containing 25 µM APV and DNXQ. D) GCaMP6s fraction of burst 1 (area under the curve) for burst 2 after 5 uM nimodipine incubation for each condition. E) GCaMP6s fraction of burst 1 (area under the curve) for burst 3 after 30 µM nimodipine incubation for each condition. Bars depict mean ± SEM. each dot represents an individual neuron. E) Boxplot of DCV fusion events per field of view in mass cultured neurons during burst 1 relative to fusion in control hippocampal neurons (Figure 1C) grown on glial micro-islands. Boxplots represent median (line), mean (+) and Tukey range (whiskers). Each dot represents an individual neuron.

## Notes

### Competing Interest Statement

The authors have declared no competing interest.

### Summary of Updates

text has been updated. The figures remain as they were.

## References

1. Altierab, C., Dubela, S. J., Barrèrea, C., Jarviscd, S. E., Stotzce, S. C., Spaetgenscf, R. L., Scottg, J. D., Corneth, V., De Waardi, M., Zamponicj, G. W., Nargeota, J., & Bourinetak, E. (2002). Trafficking of L-type calcium channels mediated by the postsynaptic scaffolding protein AKAP79. Journal of Biological Chemistry, 277(37), 33598–33603. 10.1074/jbc.M202476200

2. Arora, S., Saarloos, I., Kooistra, R., van de Bospoort, R., Verhage, M., & Toonen, R. F. (2017). SNAP-25 gene family members differentially support secretory vesicle fusion. Journal of Cell Science, 130(11), 1877–1889. 10.1242/jcs.201889

3. Balkowiec, A., & Katz, D. M. (2002). Cellular mechanisms regulating activity-dependent release of native brain-derived neurotrophic factor from hippocampal neurons. Journal of Neuroscience, 22(23), 10399–10407. 10.1523/jneurosci.22-23-10399.2002

4. Blöchl, A., & Thoenen, H. (1995). Characterization of Nerve Growth Factor (NGF) Release from Hippocampal Neurons: Evidence for a Constitutive and an Unconventional Sodium-dependent Regulated Pathway. European Journal of Neuroscience, 7(6), 1220–1228. 10.1111/J.1460-9568.1995.TB01112.X

5. Camello, C., Lomax, R., Petersen, O. H., & Tepikin, A. v. (2002). Calcium leak from intracellular store - The enigma of calcium signalling. Cell Calcium, 32(5–6), 355–361. 10.1016/S0143416002001926

6. Canossa, M., Gärtner, A., Campana, G., Inagaki, N., & Thoenen, H. (2001). Regulated secretion of neurotrophins by metabotropic glutamate group I (mGIuRI) and Trk receptor activation is mediated via phospholipase C signalling pathways. EMBO Journal, 20(7), 1640–1650. 10.1093/emboj/20.7.1640

7. Castillo, P. E., Weisskopf, M. G., & Nicoll, R. A. (1994). The role of Ca2+ channels in hippocampal mossy fiber synaptic transmission and long-term potentiation. Neuron, 12(2), 261–269. 10.1016/0896-6273(94)90269-0

8. Chanaday, N. L., Nosyreva, E., Shin, O.-H., Atasoy, D., Zhang, H., Aklan, I., Bezprozvanny, I., & Kavalali, E. T. (2021). Presynaptic store-operated Ca 2+ entry drives excitatory spontaneous neurotransmission and augments endoplasmic reticulum stress. Neuron, 109. 10.1016/j.neuron.2021.02.023

9. Chen, T.-W., Wardill, T. J., Sun, Y., Pulver, S. R., Renninger, S. L., Baohan, A., Schreiter, E. R., Kerr, R. A., Orger, M. B., Jayaraman, V., Looger, L. L., Svoboda, K., & Kim, D. S. (2013). Ultrasensitive fluorescent proteins for imaging neuronal activity. 10.1038/nature12354

10. Chen, Z., Pan, S., Yin, K., Zhang, Y., Yuan, X., Wang, S., Yang, S., Shen, Q., Tang, Y., Li, J., Wang, Y., Lu, Y., & Zhang, G. (2021). Deficiency of ER Ca2+ sensor STIM1 in AgRP neurons confers protection against dietary obesity. Cell Reports, 37(3), 109868. 10.1016/J.CELREP.2021.109868

11. Cheng, P. L., Song, A. H., Wong, Y. H., Wang, S., Zhang, X., & Poo, M. M. (2011). Self-amplifying autocrine actions of BDNF in axon development. Proceedings of the National Academy of Sciences of the United States of America, 108(45), 18430–18435. 10.1073/pnas.1115907108

12. de Juan-Sanz, J., Holt, G. T., Schreiter, E. R., de Juan, F., Kim, D. S., & Ryan, T. A. (2017). Axonal Endoplasmic Reticulum Ca2+ Content Controls Release Probability in CNS Nerve Terminals. Neuron, 93(4), 867–881.e6. 10.1016/j.neuron.2017.01.010

13. De Wit, J., Toonen, R. F., & Verhage, M. (2009). Matrix-dependent local retention of secretory vesicle cargo in cortical neurons. Journal of Neuroscience, 29(1), 23–37. 10.1523/JNEUROSCI.3931-08.2009

14. Dickson, E. J., Duman, J. G., Moody, M. W., Chen, L., & Hille, B. (2012). Orai-STIM-mediated Ca2+ release from secretory granules revealed by a targeted Ca2+ and pH probe. Proceedings of the National Academy of Sciences of the United States of America, 109(51), E3539. 10.1073/PNAS.1218247109/-/DCSUPPLEMENTAL

15. Dittmer, P. J., Dell’Acqua, M. L., & Sather, W. A. (2019). Synaptic crosstalk conferred by a zone of differentially regulated Ca2+ signaling in the dendritic shaft adjoining a potentiated spine. Proceedings of the National Academy of Sciences of the United States of America, 116(27), 13611–13620. 10.1073/PNAS.1902461116/-/DCSUPPLEMENTAL

16. Dittmer, P. J., Wild, A. R., Dell’Acqua, M. L., & Sather, W. A. (2017). STIM1 Ca2+ Sensor Control of L-type Ca2+-Channel-Dependent Dendritic Spine Structural Plasticity and Nuclear Signaling. Cell Reports, 19(2), 321–334. 10.1016/j.celrep.2017.03.056

17. Dolmetsch, R. E., Pajvani, U., Fife, K., Spotts, J. M., & Greenberg, M. E. (2001). Signaling to the nucleus by an L-type calcium channel- calmodulin complex through the MAP kinase pathway. Science, 294(5541), 333–339. 10.1126/science.1063395

18. Emptage, N. J., Reid, C. A., & Fine, A. (2001). Calcium Stores in Hippocampal Synaptic Boutons Mediate Short-Term Plasticity, Store-Operated Ca2+ Entry, and Spontaneous Transmitter Release. Neuron, 29(1), 197–208. 10.1016/S0896-6273(01)00190-8

19. Farina, M., van de Bospoort, R., He, E., Persoon, C. M., van Weering, J. R. T., Broeke, J. H., Verhage, M., & Toonen, R. F. (2015a). CAPS-1 promotes fusion competence of stationary dense-core vesicles in presynaptic terminals of mammalian neurons. ELife, 2015(4). 10.7554/eLife.05438

20. Farina, M., van de Bospoort, R., He, E., Persoon, C. M., van Weering, J. R. T., Broeke, J. H., Verhage, M., & Toonen, R. F. (2015b). CAPS-1 promotes fusion competence of stationary dense-core vesicles in presynaptic terminals of mammalian neurons. ELife, 2015(4). 10.7554/eLife.05438

21. García, A. G., García-De-Diego, A. M., Gandía, L., Borges, R., & García-Sancho, J. (2006). Calcium signaling and exocytosis in adrenal chromaffin cells. Physiological Reviews, 86(4), 1093–1131. 10.1152/PHYSREV.00039.2005/ASSET/IMAGES/LARGE/Z9J00406241 30010.JPEG

22. Gärtner, A., & Staiger, V. (2002). Neurotrophin secretion from hippocampal neurons evoked by long-term-potentiation-inducing electrical stimulation patterns. Proceedings of the National Academy of Sciences of the United States of America, 99(9), 6386–6391. 10.1073/pnas.092129699

23. Granseth, B., Odermatt, B., Royle, S. J. J., & Lagnado, L. (2006). Clathrin-Mediated Endocytosis Is the Dominant Mechanism of Vesicle Retrieval at Hippocampal Synapses. Neuron, 51(6), 773–786. 10.1016/j.neuron.2006.08.029

24. Griesbeck, O., Canossa, M., Campana, G., Gärtner, A., Hoener, M. C., Nawa, H., Kolbeck, R., & Thoenen, H. (1999). Are there differences between the secretion characteristics of NGF and BDNF? Implications for the modulatory role of neurotrophins in activity-dependent neuronal plasticity. Microscopy Research and Technique, 45(4–5), 262–275. 10.1002/(SICI)1097-0029(19990515/01)45:4/5<262::AID-JEMT10>3.0.CO;2-K

25. He, M., Bodi, I., Mikala, G., & Schwartz, A. (1997). Motif III S5 of L-type Calcium Channels Is Involved in the Dihydropyridine Binding Site. Journal of Biological Chemistry, 272(5), 2629–2633. 10.1074/jbc.272.5.2629

26. Hinckelmann, M. V., Virlogeux, A., Niehage, C., Poujol, C., Choquet, D., Hoflack, B., Zala, D., & Saudou, F. (2016). Self-propelling vesicles define glycolysis as the minimal energy machinery for neuronal transport. Nature Communications 2016 *7*:1, *7*(1), 1–13. 10.1038/ncomms13233

27. Iwasaki, S., Momiyama, A., Uchitel, O. D., & Takahashi, T. (2000). Developmental Changes in Calcium Channel Types Mediating Central Synaptic Transmission. Journal of Neuroscience, 20(1), 59–65. 10.1523/JNEUROSCI.20-01-00059.2000

28. JoséGarrido, J., Giraud, P., Carlier, E., Fernandes, F., Moussif, A., Fache, M. P., Debanne, D., & Dargent, B. (2003). A targeting motif involved in sodium channel clustering at the axonal initial segment. Science, 300(5628), 2091–2094. 10.1126/SCIENCE.1085167/SUPPL_FILE/GARRIDO.SOM.PDF

29. Kamijo, S., Ishii, Y., Horigane, S. I., Suzuki, K., Ohkura, M., Nakai, J., Fujii, H., Takemoto-Kimura, S., & Bito, H. (2018). A Critical Neurodevelopmental Role for L-Type Voltage-Gated Calcium Channels in Neurite Extension and Radial Migration. Journal of Neuroscience, 38(24), 5551–5566. 10.1523/JNEUROSCI.2357-17.2018

30. Kolarow, R., Brigadski, T., & Lessmann, V. (2007). Postsynaptic secretion of BDNF and NT-3 from hippocampal neurons depends on calcium-calmodulin kinase II signaling and proceeds via delayed fusion pore opening. Journal of Neuroscience, 27(39), 10350– 10364. 10.1523/JNEUROSCI.0692-07.2007

31. Leitch, B., Szostek, A., Lin, R., & Shevtsova, O. (2009). Subcellular distribution of L-type calcium channel subtypes in rat hippocampal neurons. Neuroscience, 164(2), 641–657. 10.1016/J.NEUROSCIENCE.2009.08.006

32. Llano, I., González, J., Caputo, C., Lai, F. A., Blayney, L. M., Tan, Y. P., & Marty, A. (2000). Presynaptic calcium stores underlie large-amplitude miniature IPSCs and spontaneous calcium transients. Nature Neuroscience 2000 *3*:12, *3*(12), 1256–1265. 10.1038/81781

33. Luarte, A., Cornejo, V. H., Bertin, F., Gallardo, J., & Couve, A. (2018). The axonal endoplasmic reticulum: One organelle—many functions in development, maintenance, and plasticity. Developmental Neurobiology, 78(3), 181–208. http://doi.wiley.com/10.1002/dneu.22560

34. Luebke, J. I., Dunlap, K., & Turner, T. J. (1993). Multiple calcium channel types control glutamatergic synaptic transmission in the hippocampus. Neuron, 11(5), 895–902. 10.1016/0896-6273(93)90119-C

34. Lundberg, J. M., Rudehill, A., Sollevi, A., Theodorsson-Norheim, E., & Hamberger, B. (1986). Frequency- and reserpine-dependent chemical coding of sympathetic transmission: Differential release of noradrenaline and neuropeptide Y from pig spleen. Neuroscience Letters, 63(1), 96–100. 10.1016/0304-3940(86)90020-0

35. Malva, J. O., Xapelli, S., Baptista, S., Valero, J., Agasse, F., Ferreira, R., & Silva, A. P. (2012). Multifaces of neuropeptide Y in the brain – Neuroprotection, neurogenesis and neuroinflammation. Neuropeptides, 46(6), 299–308. 10.1016/J.NPEP.2012.09.001

36. Martín, E. D., & Buño, W. (2003). Caffeine-mediated presynaptic long-term potentiation in hippocampal CA1 pyramidal neurons. Journal of Neurophysiology, 89(6), 3029–3038. 10.1152/JN.00601.2002/ASSET/IMAGES/LARGE/9K0633126007.JPEG

37. Mcpherson, P. S., Kim, Y.-K., Valdivia, H., Knudson, + C Michael, Takekura, H., Franzini-Armstrong, C., Coronado, R., Campbell, K. P., Program, *, & Neuroscience, I. (1991). The Brain Ryanodine Receptor: A Caffeine-Sensitive Calcium Release Channel. Neuron, 7, 17–25.

38. Mennerick, S., Que, J., Benz, A., & Zorumski, C. F. (1995). Passive and synaptic properties of hippocampal neurons grown in microcultures and in mass cultures. Journal of Neurophysiology, 73(1), 320–332. 10.1152/jn.1995.73.1.320

39. Meyer-Lindenberg, A., Domes, G., Kirsch, P., & Heinrichs, M. (2011). Oxytocin and vasopressin in the human brain: social neuropeptides for translational medicine. Nature Reviews. Neuroscience, 12(9), 524–538. 10.1038/NRN3044

40. Moro, A., Hoogstraaten, R. I., Persoon, C. M., Verhage, M., & Toonen, R. F. (2021). Quantitative analysis of dense-core vesicle fusion in rodent CNS neurons. STAR Protocols, 2(1), 100325. 10.1016/J.XPRO.2021.100325

41. Moro, A., van Woerden, G. M., Toonen, R. F., & Verhage, M. (2020). CaMKII controls neuromodulation via neuropeptide gene expression and axonal targeting of neuropeptide vesicles. PLoS Biology, 18(8), e3000826. 10.1371/journal.pbio.3000826

42. Mukherjee, M., Mukherjee, C., Ghosh, V., Jain, A., Sadhukhan, S., Dagar, S., & Sahu, B. S. (2022). ER Stress Impedes Regulated Secretion by Governing Key Granulogenic and Exocytotic Molecular Switches. SSRN Electronic Journal, 2023.04.18.537291. 10.2139/ssrn.4285379

43. Nagai, T., Ibata, K., Park, E. S., Kubota, M., Mikoshiba, K., & Miyawaki, A. (2002). A variant of yellow fluorescent protein with fast and efficient maturation for cell-biological applications. Nature Biotechnology, 20(1), 87–90. 10.1038/nbt0102-87

44. Nakamura, Y., Harada, H., Kamasawa, N., Matsui, K., Rothman, J. S., Shigemoto, R., Silver, R. A., DiGregorio, D. A., & Takahashi, T. (2015). Nanoscale Distribution of Presynaptic Ca2+ Channels and Its Impact on Vesicular Release during Development. Neuron, 85(1), 145–158. 10.1016/J.NEURON.2014.11.019

45. Naldini, L., Blömer, U., Gallay, P., Ory, D., Mulligan, R., Gage, F. H., Verma, I. M., & Trono, D. (1996). In vivo gene delivery and stable transduction of nondividing cells by a lentiviral vector. Science, 272(5259), 263–267. 10.1126/science.272.5259.263

46. Pang, P. T., Teng, H. K., Zaitsev, E., Woo, N. T., Sakata, K., Zhen, S., Teng, K. K., Yung, W. H., Hempstead, B. L., & Lu, B. (2004). Cleavage of proBDNF by tPA/plasmin is essential for long-term hippocampal plasticity. Science, 306(5695), 487–491. 10.1126/SCIENCE.1100135/SUPPL_FILE/PANG.SOM.PDF

47. Panzera, L. C., Johnson, B., Quinn, J. A., Cho, I. H., Tamkun, M. M., & Hoppa, M. B. (2022). Activity-dependent endoplasmic reticulum Ca2+ uptake depends on Kv2.1-mediated endoplasmic reticulum/plasma membrane junctions to promote synaptic transmission. Proceedings of the National Academy of Sciences of the United States of America, 119(30). 10.1073/PNAS.2117135119/-/DCSUPPLEMENTAL

48. Park, C. Y., Shcheglovitov, A., & Dolmetsch, R. (2010). The CRAC channel activator STIM1 binds and inhibits L-type voltage-gated calcium channels. Science, 330(6000), 101–105. 10.1126/science.1191027

49. Pavez, M., Thompson, A. C., Arnott, H. J., Mitchell, C. B., D’Atri, I., Don, E. K., Chilton, J. K., Scott, E. K., Lin, J. Y., Young, K. M., Gasperini, R. J., & Foa, L. (2019). STIM1 is required for remodeling of the endoplasmic reticulum and microtubule cytoskeleton in steering growth cones. Journal of Neuroscience, 39(26), 5095–5114. 10.1523/JNEUROSCI.2496-18.2019

50. Persoon, C. M., Hoogstraaten, R. I., Nassal, J. P., van Weering, J. R. T., Kaeser, P. S., Toonen, R. F., Verhage, M., Persoon, Claudia M, Hoogstraaten, Rein I, Nassal, Jorris P, van Weering, Jan RT, Kaeser, Pascal S, Toonen, Ruud F, Verhage, M., Persoon, C. M., Hoogstraaten, R. I., Nassal, J. P., van Weering, J. R. T., Kaeser, P. S., Toonen, R. F., & Verhage, M. (2019). The RAB3-RIM Pathway Is Essential for the Release of Neuromodulators. Neuron, 104(6), 1065–1080.e12. 10.1016/j.neuron.2019.09.015

51. Persoon, C. M., Moro, A., Nassal, J. P., Farina, M., Broeke, J. H., Arora, S., Dominguez, N., Weering, J. R., Toonen, R. F., & Verhage, M. (2018). Pool size estimations for dense-core vesicles in mammalian CNS neurons . The EMBO Journal, 37(20), e99672. 10.15252/embj.201899672

52. Puntman, D. C., Arora, S., Farina, M., Toonen, R. F., & Verhage, M. (2021). Munc18-1 Is Essential for Neuropeptide Secretion in Neurons. Journal of Neuroscience, 41(28), 5980–5993. 10.1523/JNEUROSCI.3150-20.2021

53. Rao, W., Zhang, L., Peng, C., Hui, H., Wang, K., Su, N., Wang, L., Dai, S. hui, Yang, Y. fan, Chen, T., Luo, P., & Fei, Z. (2015). Downregulation of STIM2 improves neuronal survival after traumatic brain injury by alleviating calcium overload and mitochondrial dysfunction. Biochimica et Biophysica Acta (BBA) - Molecular Basis of Disease, 1852(11), 2402– 2413. 10.1016/J.BBADIS.2015.08.014

54. Schmidt, E. K., Clavarino, G., Ceppi, M., & Pierre, P. (2009). SUnSET, a nonradioactive method to monitor protein synthesis. 10.1038/NMETH.1314

55. Scholz, K. P., & Miller, R. J. (1995). Developmental changes in presynaptic calcium channels coupled to glutamate release in cultured rat hippocampal neurons. Journal of Neuroscience, 15(6), 4612–4617. 10.1523/JNEUROSCI.15-06-04612.1995

56. Shakiryanova, D., Tully, A., Hewes, R. S., Deitcher, D. L., & Levitan, E. S. (2005). Activity-dependent liberation of synaptic neuropeptide vesicles. Nature Neuroscience 2005 *8*:2, *8*(2), 173–178. 10.1038/nn1377

57. Shakiryanova, D., Zettel, G. M., Gu, T., Hewes, R. S., & Levitan, E. S. (2011). Synaptic neuropeptide release induced by octopamine without Ca2+ entry into the nerve terminal. Proceedings of the National Academy of Sciences of the United States of America, 108(11), 4477–4481. 10.1073/pnas.1017837108

58. Simmons, M. L., Terman, G. W., Gibbs, S. M., & Chavkin, C. (1995). L-type calcium channels mediate dynorphin neuropeptide release from dendrites but not axons of hippocampal granule cells. Neuron, 14(6), 1265–1272. 10.1016/0896-6273(95)90273-2

59. Sun, Y., Zhang, H., Selvaraj, S., Sukumaran, P., Lei, S., Birnbaumer, L., & Singh, B. B. (2017). Inhibition of L-Type Ca2+ Channels by TRPC1-STIM1 Complex Is Essential for the Protection of Dopaminergic Neurons. Journal of Neuroscience, 37(12), 3364–3377. 10.1523/JNEUROSCI.3010-16.2017

60. Takahashi, T., & Momiyama, A. (1993). Different types of calcium channels mediate central synaptic transmission. Nature, 366(6451), 156–158. 10.1038/366156A0

61. Terasaki, M. (2018). Axonal endoplasmic reticulum is very narrow. Journal of Cell Science, 131(4). 10.1242/JCS.210450/VIDEO-7

62. Tippens, A. L., Pare, J.-F., Langwieser, N., Moosmang, S., Milner, T. A., Smith, Y., & Lee, A. (2008). Ultrastructural evidence for pre- and postsynaptic localization of Cav1.2 L-type Ca2+ channels in the rat hippocampus. The Journal of Comparative Neurology, 506(4), 569–583. 10.1002/cne.21567

63. Unni, V. K., Zakharenko, S. S., Zablow, L., DeCostanzo, A. J., & Siegelbaum, S. A. (2004). Calcium Release from Presynaptic Ryanodine-Sensitive Stores Is Required for Long-Term Depression at Hippocampal CA3-CA3 Pyramidal Neuron Synapses. The Journal of Neuroscience, 24(43), 9612. 10.1523/JNEUROSCI.5583-03.2004

64. van de Bospoort, R., Farina, M., Schmitz, S. K., de Jong, A., de Wit, H., Verhage, M., & Toonen, R. F. (2012). Munc13 controls the location and efficiency of dense-core vesicle release in neurons. Journal of Cell Biology, 199(6), 883–891. 10.1083/jcb.201208024

65. Verhage, M., McMahon, H. T., Ghijsen, W. E. J. M., Boomsma, F., Scholten, G., Wiegant, V. M., & Nicholls, D. G. (1991). Differential release of amino acids, neuropeptides, and catecholamines from isolated nerve terminals. Neuron, 6(4), 517–524. 10.1016/0896-6273(91)90054-4

66. Verkhratsky, A. (2002). The endoplasmic reticulum and neuronal calcium signalling. Cell Calcium, 32(5–6), 393–404. 10.1016/S0143416002001896

67. Verkhratsky, A. (2005). Physiology and pathophysiology of the calcium store in the endoplasmic reticulum of neurons. In Physiological Reviews (Vol. 85, Issue 1, pp. 201– 279). American Physiological Society. 10.1152/physrev.00004.2004

68. Wang, Y., Deng, X., Mancarella, S., Hendron, E., Eguchi, S., Soboloff, J., Tang, X. D., & Gill, D. L. (2010). The calcium store sensor, STIM1, reciprocally controls Orai and Ca V1.2 channels. Science, 330(6000), 105–109. 10.1126/SCIENCE.1191086/SUPPL_FILE/WANG_SOM.PDF

69. Wheeler, D. B., Randall, A., & Tsien, R. W. (1994). Roles of N-Type and Q-Type Ca$^{2+}$ Channels in Supporting Hippocampal Synaptic Transmission. In New Series (Vol. 264, Issue 5155). American Association for the Advancement of Science . 10.1126/SCIENCE.7832825

70. Wierda, K. D. B., Toonen, R. F. G., de Wit, H., Brussaard, A. B., & Verhage, M. (2007). Interdependence of PKC-Dependent and PKC-Independent Pathways for Presynaptic Plasticity. Neuron, 54(2), 275–290. 10.1016/j.neuron.2007.04.001

71. Wong, M. Y., Shakiryanova, D., & Levitan, E. S. (2009). Presynaptic Ryanodine Receptor- CamKII Signaling is Required for Activity-dependent Capture of Transiting Vesicles. Journal of Molecular Neuroscience : MN, 37(2), 146. 10.1007/S12031-008-9080-8

72. Wu, Y., Whiteus, C., Xu, C. S., Hayworth, K. J., Weinberg, R. J., Hess, H. F., & de Camilli, P. (2017). Contacts between the endoplasmic reticulum and other membranes in neurons. Proceedings of the National Academy of Sciences of the United States of America, 114(24), E4859–E4867. 10.1073/PNAS.1701078114/-/DCSUPPLEMENTAL

73. Xia, X., Lessmann, V., & Martin, T. F. J. (2009). Imaging of evoked dense-core-vesicle exocytosis in hippocampal neurons reveals long latencies and kiss-and-run fusion events. Journal of Cell Science, 122(1), 75–82. 10.1242/JCS.034603

74. Yalçın, B., Zhao, L., Stofanko, M., O’Sullivan, N. C., Kang, Z. H., Roost, A., Thomas, M. R., Zaessinger, S., Blard, O., Patto, A. L., Sohail, A., Baena, V., Terasaki, M., & O’Kane, C. J. (2017). Modeling of axonal endoplasmic reticulum network by spastic paraplegia proteins. ELife, 6. 10.7554/ELIFE.23882

75. Yoshimura, H. (2005). The Potential of Caffeine for Functional Modification from Cortical Synapses to Neuron Networks in the Brain. Current Neuropharmacology, 3(4), 309. 10.2174/157015905774322543

76. Zakharenko, S. S., Patterson, S. L., Dragatsis, I., Zeitlin, S. O., Siegelbaum, S. A., Kandel, E. R., & Morozov, A. (2003). Presynaptic BDNF Required for a Presynaptic but Not Postsynaptic Component of LTP at Hippocampal CA1-CA3 Synapses. Neuron, 39(6), 975–990. 10.1016/S0896-6273(03)00543-9

77. Zhang, H., Fu, Y., Altier, C., Platzer, J., Surmeier, D. J., & Bezprozvanny, I. (2006). CaV1.2 and CaV1.3 neuronal L-type calcium channels: Differential targeting and signaling to pCREB. European Journal of Neuroscience, 23(9), 2297–2310. 10.1111/J.1460-9568.2006.04734.X

78. Zhang, H., Maximov, A., Fu, Y., Xu, F., Tang, T. S., Tkatch, T., Surmeier, D. J., & Bezprozvanny, I. (2005). Association of CaV1.3 L-Type Calcium Channels with Shank. Journal of Neuroscience, 25(5), 1037–1049. 10.1523/JNEUROSCI.4554-04.2005

